# Combined, yet Separate: cocktails of carriers (not drugs) for α-particle therapy of solid tumors expressing moderate-to-low levels of targetable markers

**DOI:** 10.1101/2023.07.31.551152

**Authors:** Rajiv Ranjit Nair, Aprameya Prasad, Omkar Bhatavdekar, Aira Sarkar, Kathleen L. Gabrielson, Stavroula Sofou

## Abstract

Alpha-particle radionuclide-antibody conjugates are being clinically evaluated against solid cancers expressing moderate levels of the targeted markers, with promising results. These findings are attributed to the high killing power of alpha-particles in spite of the expected decrease in antibody tumor uptake, that reduces tumor absorbed doses. However, when tumor absorbed doses are reduced, addressing the heterogeneities in delivery of alpha-particles within solid tumors (i.e. enabling uniform irradiation patterns) becomes critical: to maintain efficacy, the fewer alpha-particles delivered within tumors need to traverse/hit as many different cancer cells as possible. This proof-of-concept study describes an approach to complement the antibody-targeted radiotherapy by using a separate carrier to deliver a fraction of the injected radioactivity to tumor regions geographically different than those affected by the antibody; collectively, the two carriers should distribute the alpha-particle emitters, Actinium-225 in particular, more uniformly within tumors maintaining efficacy.

**Methods:** We monitored the extent(s) of tumor growth inhibition, onset delay of spontaneous metastases and/or survival on orthotopic MDA-MB-213 and MDA-MB-436 triple negative breast cancer mouse models and on an ectopic BxPC3 pancreatic cancer mouse model, treated systemically with the two separate carriers. Tumors were chosen to express different (but low) levels of HER1, utilized as a model antibody-targeted marker.

**Results:** Independent of tumor origin and/or resistance to chemotherapy, the two separate carriers: (a) improved the ‘primary’ tumor growth inhibition, (b) eliminated the formation of spontaneous metastases, and/or (c) prolonged survival, at lower or comparable tumor delivered doses relative to the antibody alone, without noticeable off-target toxicities.

**Conclusion:** This tumor-agnostic strategy is timely and could be used to enhance the efficacy of existing alpha-particle radionuclide-antibody treatments without increasing, possibly even reducing, the total administered radioactivity.

## Introduction

For short-range, locally acting radiotherapeutic agents, the spatiotemporal microdistributions of delivered radioactivity within solid tumors could strongly affect the extent of tumor growth inhibition and treatment outcomes (*1–3*). High energy (1-10 MeV), short range (40-100μm in tissue) alpha particles (α-particles) are ideal tools to demonstrate the role of intratumoral drug microdistributions in tumor growth control. Because of their major killing mechanism, which relies on double-strand DNA breaks, α-particle irradiation is essentially impervious to cell resistance and largely independent of the tumor microenvironment (*4*). Traditionally, tumor response has been correlated with the tumor absorbed dose, that is the delivered energy averaged *over the entire tumor mass* (*5*). However, in α-particle therapy, the microdistributions of delivered radioactivity within solid tumors also matter, since cells not being hit by α-particles may likely not be killed (*2,6*).

Accordingly, for vascularized soft-tissue tumors, we previously showed that a higher tumor-absorbed dose delivered in a heterogeneous manner by, for example, employing a strongly binding/internalizing radiolabeled targeting antibody, which was mostly localized only within a certain distance from the neovasculature, was *less effective* in inhibiting tumor growth than a lower tumor-absorbed dose that was more widely spread over the entire tumor (*1*). The latter approach was enabled by splitting the same total injected radioactivity between the targeting antibodies and separately administered tumor-responsive liposomes, delivering their radioactive cargo *to complementary areas of the same tumor*. This approach was proven in different tumor xenografts formed by cancer cells overexpressing known targeted receptors (3+ score by immunohistochemistry, IHC, or > 1,000,000 receptor copies per cancer cell (*7*)) (*1,8*).

However, most cancers do not necessarily overexpress known tumor-associated antigens (*9,10*). Currently, in the clinic, antibody-conjugates with α-particle emitters are evaluated against solid tumors with low marker expression (NCT04147819). In these latter cases, not only the patterns of tumor irradiation by α-particles will still be non-uniform (*8,11*), but the tumor absorbed doses are expected to be lower than the absorbed doses of tumors with greater surface marker expression; this scenario would possibly compromise the therapeutic efficacy of the delivered α- particle therapy. Therefore, there is a need to *improve the intratumoral spreading* of α-particle emitters on solid tumors expressing moderate to low levels of targeted markers to compensate for the relatively lower delivered doses by targeting radionuclide-antibody conjugates.

In the present study, the goal was two-fold. First, to demonstrate that a treatment strategy that combines separate carriers (of the same drug), enabling complementary intratumoral drug microdistributions, can be a general delivery approach that is effective even when the expression of antibody-targeted receptors on cancer cells is moderate (<2+ by IHC; i.e. <500,000 marker copies per cell (*7*)) or even low (1+ by IHC, i.e. < 200,000 marker copies per cell (*7*)). Second, because of the applicability of this delivery strategy against cancers with receptors not being overexpressed, to demonstrate that tumor-antibody pairs perceived as “unjustifiable,” by established treatment schemes, can still be designed to be successful in effectively inhibiting tumor growth.

Herein, we evaluated the “two-carrier approach” for the delivery of the α-particle emitter Actinium- 225 (^225^Ac) using a HER1-targeting antibody, Cetuximab, in the following diverse mouse models: (1) moderately HER1-expressing MDA-MB-231 triple-negative breast cancer (TNBC) orthotopic xenografts that develop spontaneous metastases, (2) moderately HER1-expressing *cisplatin-resistant* MDA-MB-436 orthotopic xenografts that also develop spontaneous metastases, and (3) low HER1-expressing BxPC3 pancreatic cancer ectopic xenografts. We assessed the extent of tumor growth inhibition and the onset delay of metastatic growth (models (1) and (2)) and/or the extent of prolonging survival (model (3)) based on the same total injected radioactivity when delivered (a) by the combined carriers and (b) by each carrier alone.

We hypothesized that this approach may create new therapeutic opportunities in solid tumor treatment with α-particle emitters using existing antibodies and, possibly, relax the necessity to discover new highly expressed target(s) and/or targeting vectors.

## Materials and Methods

The materials used are described in the Supporting Information. Actinium-225 (^225^Ac), actinium chloride, were supplied by the U.S. Department of Energy Isotope Program, managed by the Office of Science for Nuclear Physics.

### Liposome formation, characterization and radiolabeling

Tumor-responsive liposomes (*8*), which were prepared using the thin-film hydration method, encapsulated DOTA or DTPA for loading with ^225^Ac or ^111^In, respectively, following an active loading method using the ionophore A23187, as previously described (*1,8*). The specific radioactivity of liposomes (after reaching secular equilibrium of Bismuth-213) was measured by counting the γ-photon emissions of Bismuth-213 at, 360-480 keV (or ^111^In at, 100-400 keV) using a γ-counter (Packard Cobra II Auto-Gamma, Model E5003).

### Antibody labeling and characterization

Antibodies were conjugated to DOTA-SCN (DTPA-SCN, or FITC-SCN) (*1,8*). For antibody radiolabeling, radioactivity (in 0.2 M HCl) was added to the chelator-conjugated antibody, and the reaction mixture was incubated at 37 °C for one hour. The radiolabeled antibody was purified using a 10DG column, and the radiochemical purity was evaluated using iTLC (*12*). Immunoreactivity of the (radio)-labelled antibody was measured by incubating the cells on ice for 1 h at a 100:1 receptor:antibody ratio and by correcting for non-specific antibody binding (*1,8*).

### Measurement of the Dissociation constant (K_D_) and the Average Antigen Expression by Cells

The binding isotherms of HER1-targeting ^111^In-DTPA-SCN-Cetuximab to cells on ice were measured, as previously reported (*1,8*). The binding curves were obtained by correcting for the antibody’s non-specific binding and for the labelled antibody’s immunoreactivity and were fit to obtain the receptor/antigen expression and K_D_.

### Cell lines and development of chemoresistance

MDA-MB-436 and MDA-MB-231 cell lines were purchased from ATCC and cultured using Dulbecco’s modified Eagle’s medium (DMEM) and Roswell Park Memorial Institute (RPMI), respectively, supplemented with 10% FBS, 100 units/mL penicillin, and 100 µg/mL streptomycin in an incubator at 37 °C and 5% CO_2_. The cell lines PANC-1 and BxPC3 were gifts from Dr. Denis Wirtz at Johns Hopkins University and were cultured in DMEM and RPMI, respectively, supplemented as above.

The <u>C</u>isplatin-<u>R</u>esistant MDA-MB-436 cell line (CR-MDA-MB-436) was developed by supplementing normal growth medium (RPMI) with increasing concentrations of cisplatin. The maintenance of CR-MDA-MB-436 resistance to cisplatin was regularly monitored throughout the study (by measurement of the IC_50_).

### Colony formation assay

Following a 6 h incubation with radioactivity, cells were washed with PBS and plated into tissue culture dishes at varying cell densities (*13*). When distinct cell colonies were observed in approximately 2 weeks (∼10 doubling times), the culture dishes were washed with water, and the colonies were fixed and stained using 6% (w/v) glutaraldehyde and 0.05% (w/v) crystal violet and counted. The survival function was evaluated as the number of treated colonies normalized to the number of untreated colonies while accounting for plating efficiency (*14*).

### Spheroid formation, spatiotemporal distributions and treatment

To form spheroids, 400 PANC-1 (used as an example of pancreatic cancer cells that form spheroids), 300 CR-MDA-MB-436, and/or 500 MDA-MB-231 cells (in Matrigel^TM^ at 2.5% v/v) were seeded per well in PolyHEMA-coated 96-well round-bottom plates and centrifuged for 10 min at 1000 rcf and 4 °C. The cell line BxPC3 did not form spheroids.

The spatiotemporal profiles of the fluorescently labelled HER1-targeting antibody, the tumor-responsive liposomes (the carrier), and their encapsulated fluorescent contents (used as a surrogate of ^225^Ac) were measured in 400 µm diameter spheroids, as described in detail in the supporting information (*8*).

In treatment studies, spheroids were incubated with 3.7 kBq/mL ^225^Ac, which was split into varying ratios of liposomes and the antibody for 6 and 24 h, respectively, to roughly match their blood clearance kinetics in mice. The antibody mass was 200 times in excess of the HER1 receptors expressed by all cells comprising the spheroid. After incubation, the treated spheroids were first transferred into fresh media (one spheroid per well in PolyHEMA-coated U-bottom plates) and monitored until the volume of untreated spheroids stopped growing. At that point, spheroids were transferred to adherent 96-well plates (one spheroid per well in cell culture-treated F-bottom plates), and when the untreated condition reached confluence, cells from each well were trypsinized and counted. The percentage of outgrowth/regrowth was evaluated as the number of cells counted in each treated condition normalized to the number of cells in the untreated condition.

### Animal study

Filter-top cages with sterile food and water were used to house the mice. Animal studies were performed in compliance with the Institutional Animal Care and Use Committee protocol (IACUC) guidelines. Four-to-six-week-old NSG (NOD scid gamma) 20 g female and male mice were purchased from JHU Breeding Facility. For breast cancer models, female mice were orthotopically inoculated into the second mammary fat pad through a small incision on the right side. The inoculum contained 500,000 CR-MDA-MB-436 or MDA-MB-231 cells suspended in 100µL serum-free medium. Once the tumors had grown to size, 8-10 animals were randomly assigned to the treatment condition. For the pancreatic cancer model, female and male NSG mice were subcutaneously inoculated with 500,000 BxPC3 cells suspended in 100µL of 50:50 v/v ratio of serum-free medium and Matrigel^TM^ and were treated when tumor volumes grew to 65-85mm^3^ or 100mm^3^. For treatment, all mouse models were intravenously administered 100µL of therapy at a total radioactivity of 2.96 kBq per 20 g animal. The animal weight and tumor volume were monitored daily. The tumor volume was measured using a digital caliper with a resolution of 0.01 mm. The ellipsoid formula (V=4 × π × α × β^2^/3, where α and β are the major and minor diameters, respectively) was used to calculate the tumor volume.

For the breast cancer models forming spontaneous metastases, at the endpoint of the study (14 days after therapy or 38–39 days after tumor inoculation for MDA-MB-231 xenografts; 16 days after therapy or 40–41 days after tumor inoculation for CR-MDA-MB-436 xenografts), animals were scanned by MRI to detect axillary lymph node (ALN) metastases. The endpoint was set based on the signs of tumor ulceration in the untreated cohort. For the pancreatic cancer model, ectopic tumor growth control and animal survival advantage were monitored, and the mice were sacrificed at the onset of symptoms associated with uncontrolled tumor growth.

Following euthanasia, animals from all models were dissected to recover the tumors and critical organs. Fixed tissues from harvested sections were processed and H&E stained for histological evaluation.

To evaluate the biodistribution of the carriers, tumor-bearing mice were intravenously administered 370 kBq ^111^In-DTPA-encapsulating tumor-responsive liposomes or ^111^In-DTPA- SCN-labeled HER1-targeting Cetuximab. At fixed time points, the animals were euthanized to excise the tumors and normal organs. The tissues were weighed and the associated radioactivity was measured using a γ-counter. Decay-corrected activities were reported as a percentage of the initial injected activity per mass of tissue (%IA/g).

### Dosimetry

Dosimetry was performed using the biodistributions of ^111^In-radiolabeled liposomes and antibodies based on the methodology described in ref. (*1,15*) using the software package 3D-RD- S, Radiopharmaceutical Imaging and Dosimetry, LLC (Rapid, Baltimore, MD). ^111^In has been confirmed as a surrogate for the parent ^225^Ac biodistributions (*16*). The calculations assumed that all α-particles and electron energy delivered by the antibody were absorbed by each source tissue as long as the emission occurred within the tissue. Regarding the radioactivity delivered by liposomes, 75% of the Bismuth-213 generated in the tumor from ^225^Ac decays was assumed to be retained in the tumor, and the remaining 25% was assumed to decay in the kidneys (*1*).

### Statistical Analysis

The results are reported as the arithmetic mean of n independent measurements ± standard deviation. One-way ANOVA and/or post hoc unpaired Student’s t-test were used to calculate significant differences in efficacy, with p-values < 0.05, considered to be significant. * indicates 0.01< p-values <0.05; ** p-values < 0.01.

## Results

### Characterization of carriers and cancer cells

Antibodies exhibited stable retention of radioactivity in media, and the tumor-responsive liposomes exhibited, per their design (*1,13*), pH-dependent release of the encapsulated radioactive contents (^225^Ac-DOTA and ^111^In-DTPA) (Table 1).

**Table 1.**
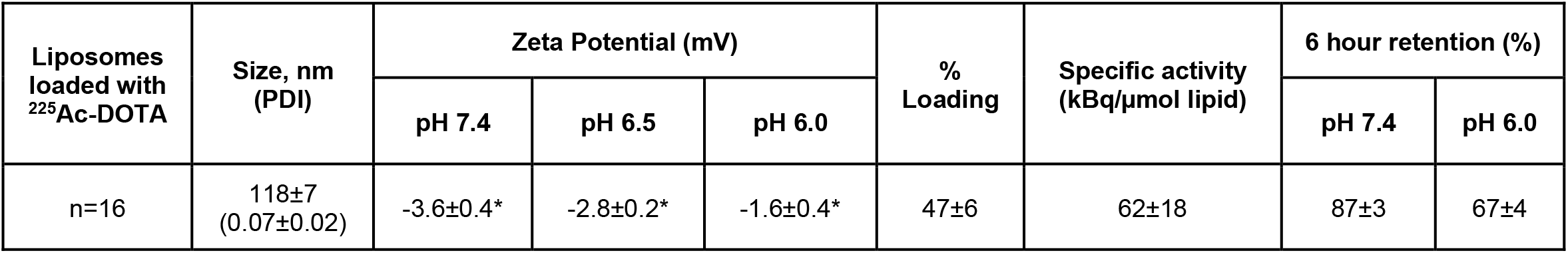
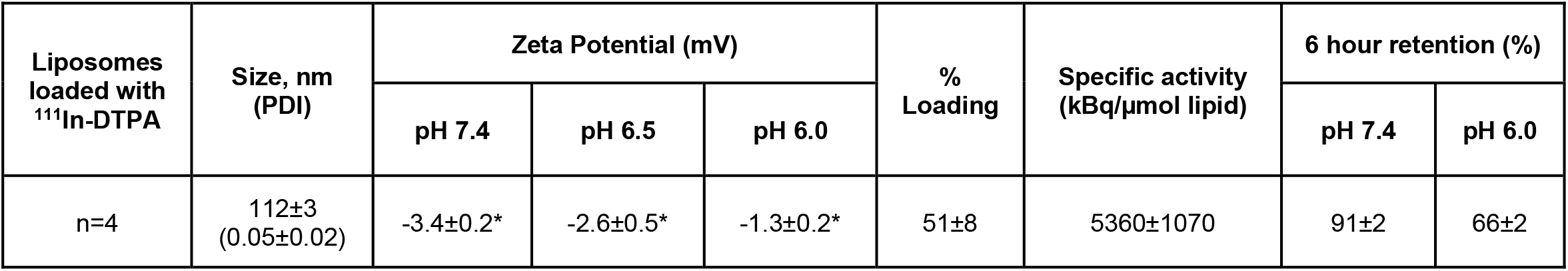
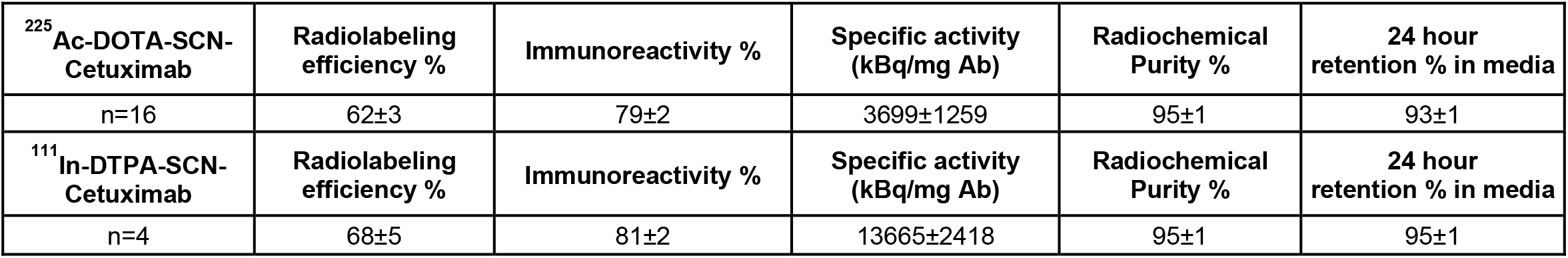
Characterization of tumor-responsive liposomes and of the HER1-targeting antibody (Cetuximab) radiolabelled with Actinium-225 and/or Indium-111. Reported are the mean values and standard deviations of repeated experiments, as indicated. * *p*-value < 0.05.

The expression levels of the HER1 receptor (Figure S1, S2 Table S1) were as follows: 520,000±120,000 in PANC1 (i.e. approximately 2+ by IHC (*17*)), 258,000±18,000 in CR-MDA-MB-436 (IHC less than 2+ but > 1+ (*17*)), 225,000±21,000 in MDA-MB-231 (IHC less than 2+ but > 1+ (*17*)), and 117,000 ±6,000 in BxPC3 (IHC 1+ (*17*)). The dissociation constant (K_D_) of radiolabeled Cetuximab across all cell lines ranged from 0.3 to 1.8 nM. Cetuximab exhibited relatively fast cell binding with extent(s) of internalization ranging from 23% in PANC1 to 32% in CR-MDA-MB-436, relative to total cell-associated antibody (Figure S3). Resistance to cisplatin of the CR-MDA-MB-436 TNBC cell line was demonstrated by the significant increase (*p*-value < 0.05) of IC_50_ relative to the wild type (1.4 ± 0.2 μg/mL to 10.2 ± 2.9 μg/mL at pH 7.4 (Figure S4a), and 1.2 ± 0.3 μg/mL to 8.2 ± 1.8 μg/mL at pH 6.0, Figure S4b).

In the absence of transport barriers, on cell monolayers, and at any given radioactivity concentration, targeted radioactivity to HER1 receptors, via ^225^Ac-DOTA-SCN-Cetixumab, resulted in lower survival compared to the non-targeted radioactivity (^225^Ac-DOTA in free or liposomal form, as expected, since liposomes were designed to not bind to cancer cells) (Figure 1a, b, d, e). The killing extent was independent of the acidification of the extracellular medium’s pH (Figure S5) that develops in the interstitium of spheroids and solid tumors (*1,8,13,18*). Importantly, the sensitivity to α-particle radiation was identical for the wild-type MDA-MB-436 cells and the cisplatin-resistant CR-MDA-MB-436 cells (Figure 1c).

**FIGURE 1.**
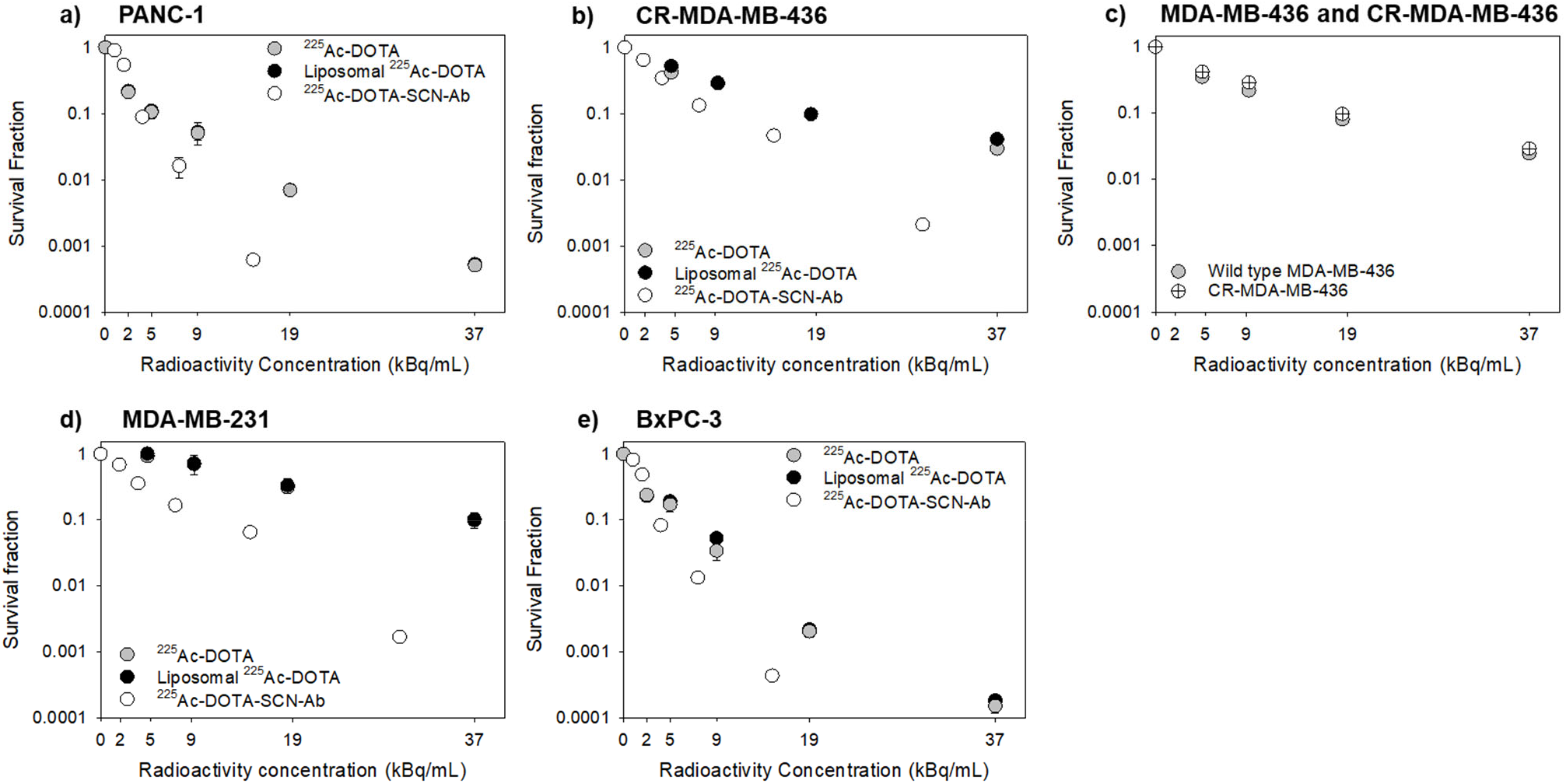
Colony survival of cancer cells following a 6-hour incubation, when in monolayers, at pH 7.4 with different radioactivity concentrations of ^225^Ac-DOTA in free form (gray circles), ^225^Ac- DOTA delivered by tumor-responsive liposomes (black circles) and the HER1-targeting ^225^Ac- DOTA-SCN-Cetuximab (white circles). Colony survival response of cisplatin-resistant CR-MDA-MB-436 cells (white crossed circles) was identical to their wild type counterparts (c), when exposed to ^225^Ac-DOTA. Data points indicate the mean values, and error bars the standard deviations of n = 3 independent experiments.

### Spheroid studies

On spheroids, contrary to studies on cell monolayers (Figure 1), the spatiotemporal microdistributions of the delivered α-particle emitters determined the level of spheroid growth inhibition. In agreement with previous studies (*1,8*), Figure 2a shows that the growth of small spheroids, with radii comparable to the range of α-particles in tissue, *was best controlled by less penetrating antibodies* (Figure 2d) because they delivered greater levels of radioactivity in spheroids (see also Figure 3A in reference (*19*)). However, outgrowth/regrowth of spheroids of larger size (Figure 2c), with radius longer than 3x the range of α-particles in tissue, was *better controlled by the tumor-responsive liposomes* (as indicated by the black arrow). This was due to the greater spheroid volume, toward the deep parts of the spheroids, which was irradiated by the highly diffusing ^225^Ac-DOTA, that was released from the liposomes within the interstitium (as shown in Figure 2f, by its fluorescent surrogate) (*2,8*), rather than the radioactivity carried by the diffusing antibodies.

**FIGURE 2.**
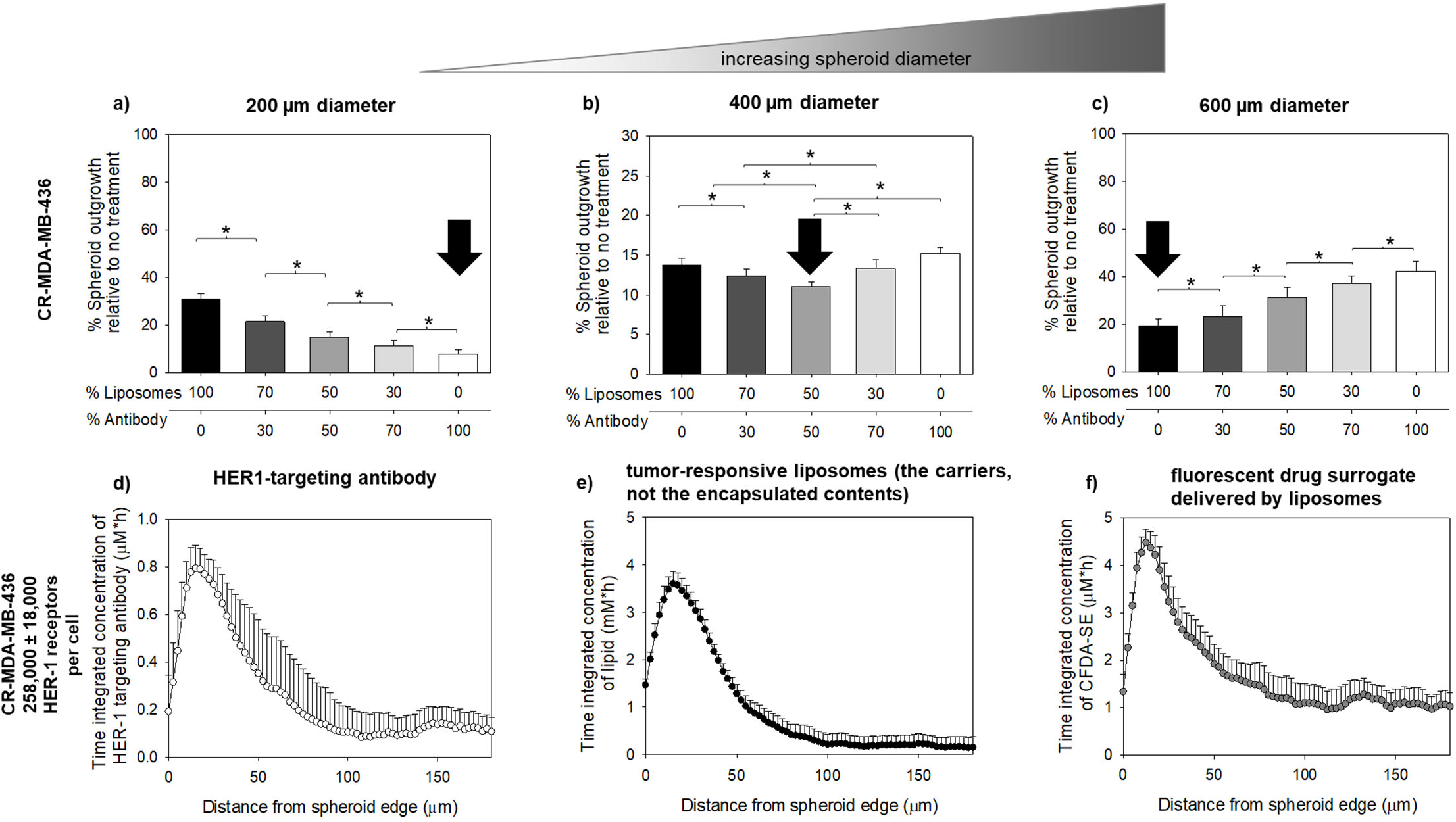
**Top panel (a-c)**. The extent of outgrowth/regrowth inhibition (used as an indirect surrogate of tumor recurrence) of CR-MDA-MB-436 spheroids of varying sizes at the time of treatment. The total radioactivity concentration was kept constant at 3.7 kBq/mL and was divided at different ratios between the tumor-responsive liposomes loaded with ^225^Ac-DOTA (incubated for 6 h) and HER1-targeting ^225^Ac-DOTA-SCN-antibody Cetuximab (incubated for 24 h). Error bars correspond to the standard deviations of repeated measurements (n=12 spheroids per condition, n=2 independent liposome and antibody preparations). **Bottom panel (d-f)**. Time-integrated spatiotemporal microdistributions of (d) the HER1-targeting antibody Cetuximab, (e) the tumor-responsive liposomes, and (f) of CFDA-SE, the fluorescent surrogate of ^225^Ac-DOTA, delivered by liposomes, in CR-MDA-MB-436 spheroids of 200µm radius. Data points indicate the mean values, and error bars the standard deviations of n = 3 different spheroids studied per time point.

**FIGURE 3.**
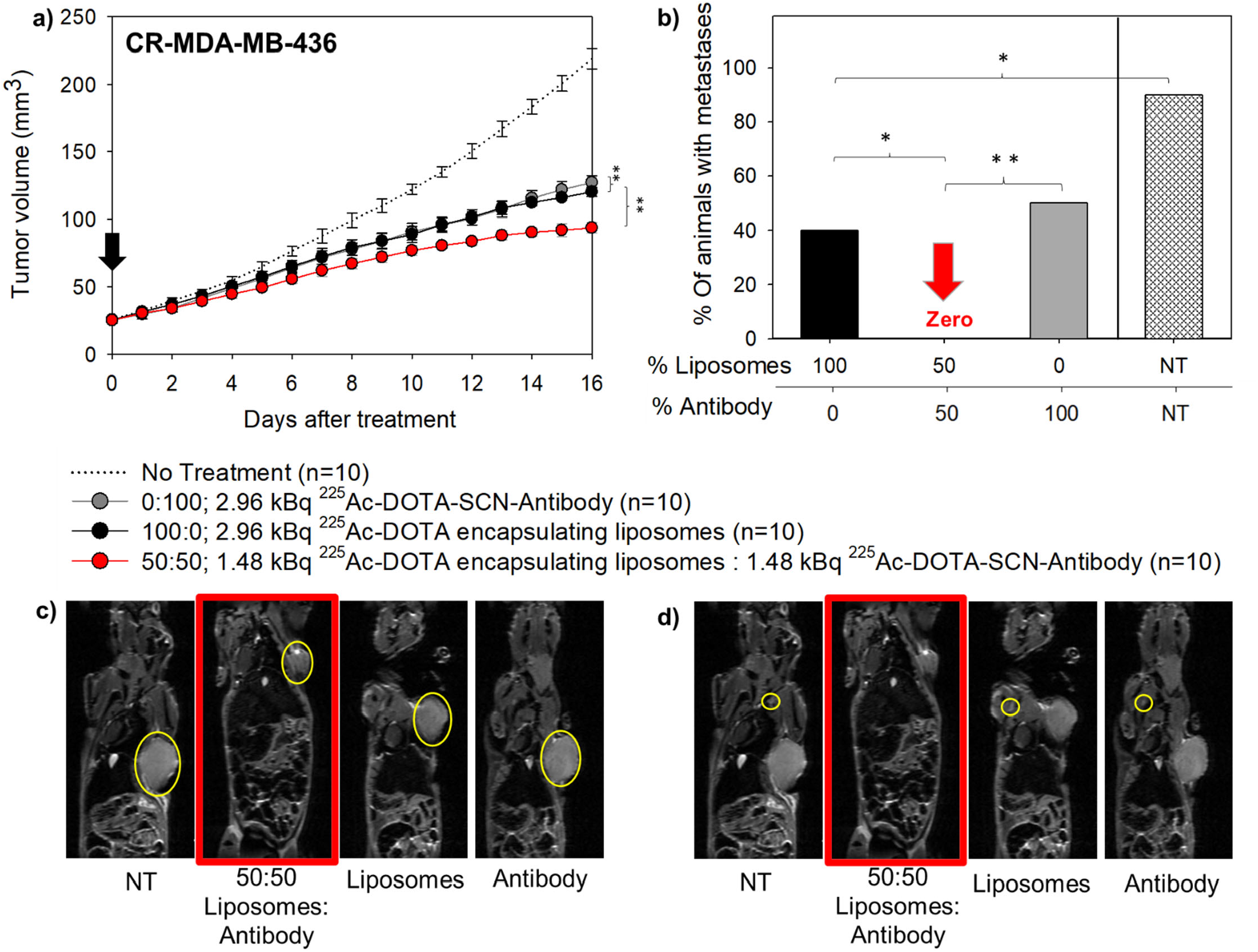
(a) Growth inhibition of primary, moderately HER1-expressing, <u>c</u>isplatin resistant CR- MDA-MB-436 orthotopic tumors, (b) onset of the spontaneous axillary lymph node (ALN) metastases, following treatment with a single i.v. injection of 2.96 kBq ^225^Ac per 20 g mouse (indicated by the black vertical arrow in (a)) delivered by: the HER1-targeting Cetuximab (grey symbols), the tumor-responsive liposomes (black symbols), and/or both, separate carriers at equal split ratio of radioactivity (red symbols). Characteristic MRI images of CR-MDA-MB-436 orthotopic tumors (c), and the ALN metastases (d) from each treatment group, acquired at the endpoint of the study. The anatomical region/level of each MRI image is focused on the frame with detectable ALN metastases. The MRI images shown serve as typical examples of animals from each cohort. Error bars correspond to the standard deviation of measurements averaged over n = 10 mice per condition as indicated. Significance was calculated with one-way ANOVA (*p*-value<0.05). Characteristic H&E-stained sections of tumors and normal organs are shown in Figure S18.

The spatiotemporal profiles of carriers and/or their (fluorescent) cargo was not affected by the origin of cancer cells forming the spheroids (Figures S6, S7). As in Figure 2, the spheroid size determined the radioactivity split ratio that best inhibited the spheroids’ outgrowth/regrowth (Figure S8).

### Dosimetry and in vivo Assessment of Efficacy

Table 2A (based on the biodistributions in Figures S9 to S12, and Tables S2 and S3) shows that only for the HER1-targeting antibody, the tumor-delivered dose decreased with decreasing HER1- expression in tumor cells (Figure S13, top panel). Normal organ dosimetry was comparable across all animal models (Table 2B).

**Table 2A.**
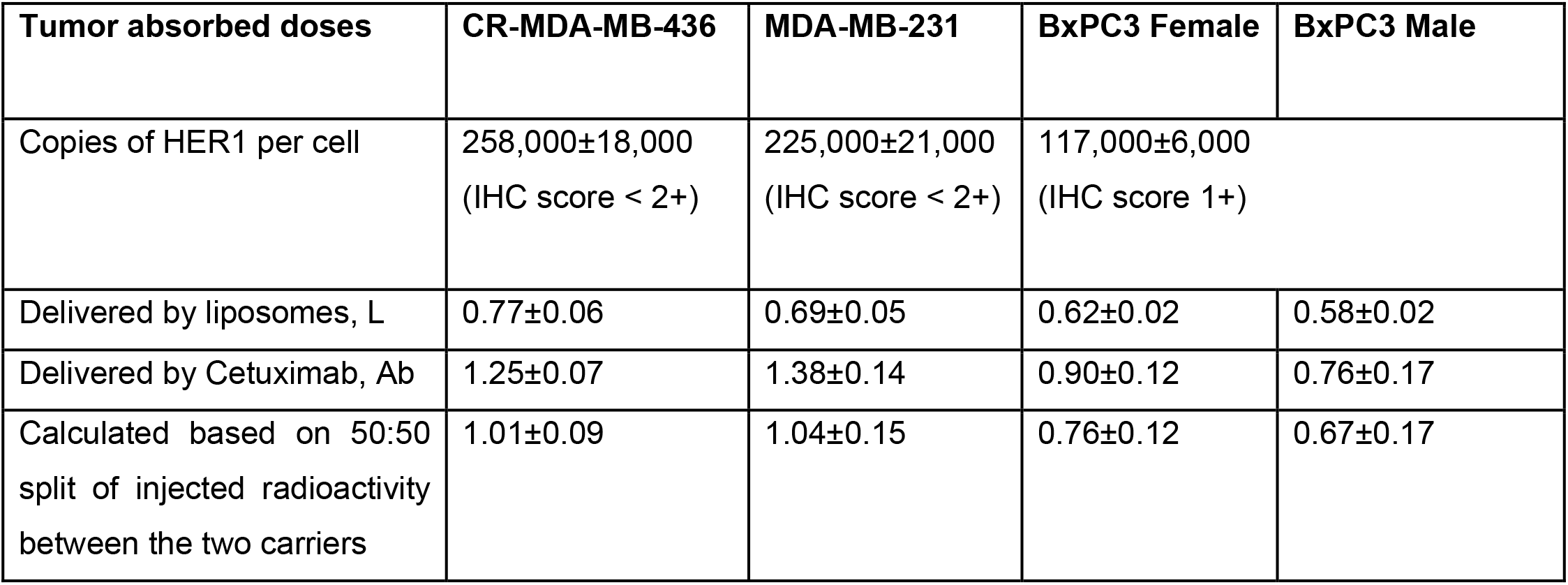
Tumor absorbed doses (Gy) from 2.96 kBq ^225^Ac delivered by each delivery modality and by their combination (at equal injected radioactivity split ratio) on the animal models studied herein. Reported are the mean values and standard deviations of measurements averaged over 3 mice per time point.

| Tumor absorbed doses | CR-MDA-MB-436 | MDA-MB-231 | BxPC3 Female | BxPC3 Male |
| --- | --- | --- | --- | --- |
| Copies of HER1 per cell | 258,000±18,000<br>(IHC score < 2+) | 225,000±21,000<br>(IHC score < 2+) | 117,000±6,000<br>(IHC score 1+) |  |
| Delivered by liposomes, L | 0.77±0.06 | 0.69±0.05 | 0.62±0.02 | 0.58±0.02 |
| Delivered by Cetuximab, Ab | 1.25±0.07 | 1.38±0.14 | 0.90±0.12 | 0.76±0.17 |
| Calculated based on 50:50 split of injected radioactivity between the two carriers | 1.01±0.09 | 1.04±0.15 | 0.76±0.12 | 0.67±0.17 |

**Table 2B.** Normal organ absorbed doses (Gy) from 2.96 kBq ^225^Ac delivered by each delivery modality on the animal models studied herein. Reported are the mean values and standard deviations of measurements averaged over 3 mice per time point.

| Absorbed Doses | CR-MDA-MB-436 |  | MDA-MB-231 |  | BxPC3 F |  | BxPC3 M |  |
| --- | --- | --- | --- | --- | --- | --- | --- | --- |
| carrier | L | Ab | L | Ab | L | Ab | L | Ab |
| heart | 0.46±0.08 | 0.79±0.12 | 0.63 ±0.05 | 1.10±0.19 | 0.35±0.03 | 1.16±0.34 | 0.33±0.06 | 0.66±0.06 |
| lungs | <0.01 | <0.01 | <0.01 | <0.01 | <0.01 | <0.01 | <0.01 | <0.01 |
| liver | 0.24±0.02 | 0.34±0.03 | 0.24±0.01 | 0.35±0.02 | 0.23±0.01 | 0.35±0.01 | 0.26±0.02 | 0.28±0.01 |
| spleen | 2.04±0.24 | 5.94±0.60 | 2.21±0.14 | 5.92±0.35 | 1.87±0.10 | 5.69±0.29 | 1.81±0.11 | 6.34±0.31 |
| kidneys | 0.51±0.22 | 0.63±0.06 | 0.49±0.05 | 0.61±0.03 | 0.32±0.07 | 0.52±0.05 | 0.44±0.05 | 0.58±0.06 |
| bone | <0.01 | <0.01 | <0.01 | <0.01 | <0.01 | <0.01 | <0.01 | <0.01 |
| muscle | <0.01 | <0.01 | <0.01 | <0.01 | <0.01 | <0.01 | <0.01 | <0.01 |

In mice with TNBC orthotopic tumors moderately expressing HER1 (IHC score < 2+), equal split of the same total injected radioactivity between the two separate carriers was accompanied by an approximately 30% decrease in the tumor absorbed dose, compared to the dose delivered by the antibody alone (Table 2A). However, the two-carrier delivery approach still resulted (1) in better growth inhibition of the orthotopic tumors (red symbols, frames, and arrows in Figures 3 and 4, panels a and c), independent of resistance to cisplatin, as demonstrated by the CR-MDA-MB-436 model, and (2) in elimination of formation of the corresponding TNBC spontaneous metastases (Figures 3 and 4, panels b and d). (Individual tumor volumes are shown in Figures S14 and S15.)

**FIGURE 4.**
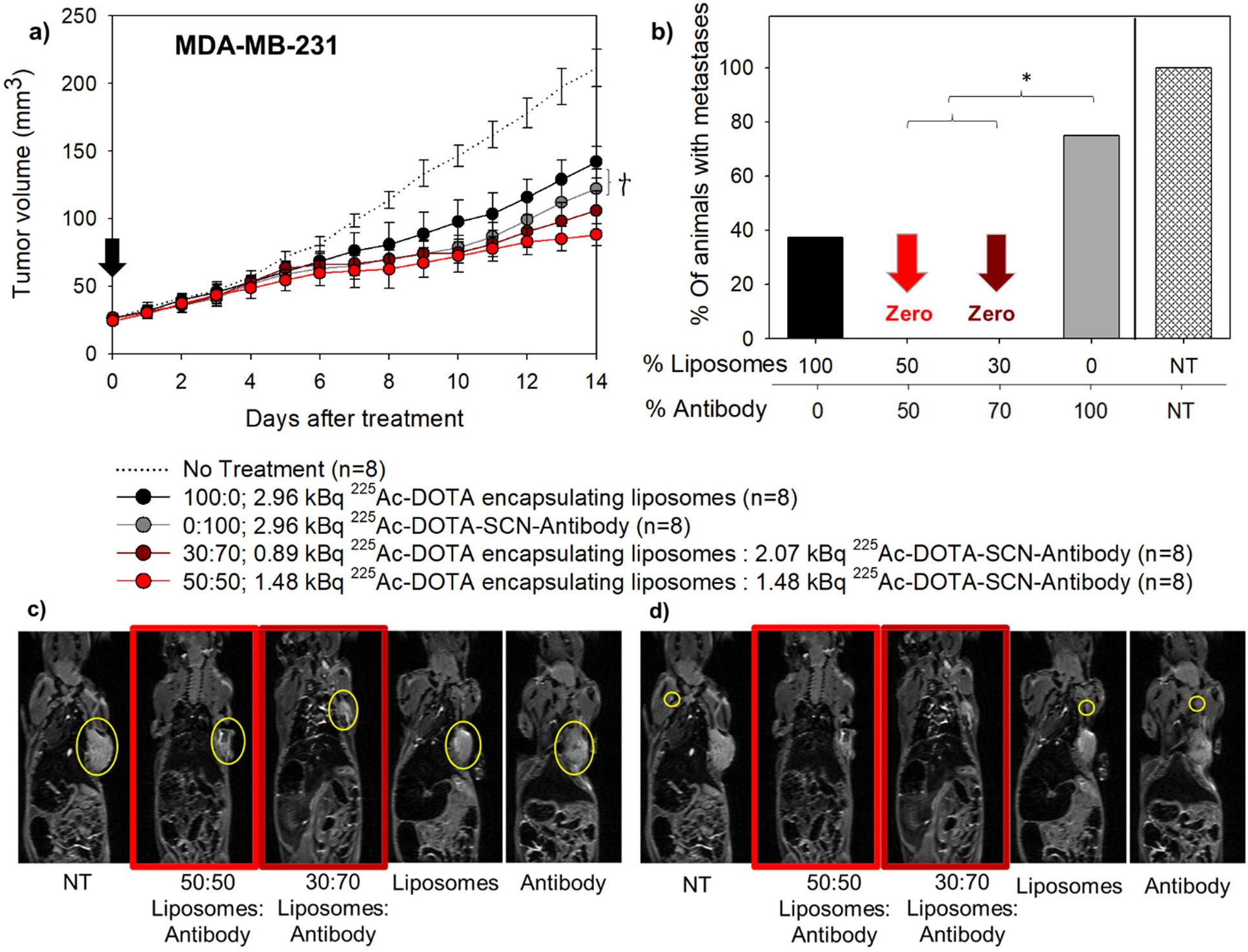
(a) Growth inhibition of primary, moderately HER1-expressing MDA-MB-231 orthotopic tumors, (b) onset delay of spontaneous axillary lymph node (ALN) metastases, following treatment with a single i.v. injection of 2.96 kBq ^225^Ac per 20 g mouse (indicated by the black vertical arrow in (a)) delivered by: the HER1-targeting Cetuximab (grey symbols), the tumor-responsive liposomes (black symbols), and/or both, separate carriers at different split ratios of radioactivity (red/dark red symbols). Characteristic MRI images of MDA-MB-213 orthotopic tumors (c), and the ALN metastases (d) from each treatment group, acquired at the endpoint of the study. Error bars correspond to the standard deviation of measurements averaged over n = 8 mice per condition as indicated. Significance was calculated with one-way ANOVA (*p*-value<0.05). In Figure a, all cohorts were statistically significant (*p*-value < 0.01) from each other except for the two groups indicated by ⴕ which were treated with activity delivered only by tumor-responsive liposomes vs. HER1-targeting Cetuximab (*p*-value=0.238). Characteristic H&E-stained sections of tumors and normal organs are shown in Figure S19.

In mice bearing subcutaneous pancreatic cancer tumors with low HER1 expression (IHC score of 1+), all delivery schemes resulted in comparable tumor-absorbed doses (Table 2A). However, the two-carrier approach best inhibited tumor growth and best prolonged survival (red symbols, lines in Figure 5, panels a, c, and b, d, respectively). Comparison of the two-carrier α-particle therapy to gemcitabine, a standard of care, again demonstrated better efficacy in terms of tumor growth control (red solid line in Figure 6a) and survival (red solid line in Figure 6b). Interestingly, survival was extended even more when the two-carrier α-particle therapy was combined with gemcitabine (red dashed line, *p*-value=0.039). On the contrary, survival was not improved when α-particle therapy delivered by each carrier alone was combined with gemcitabine (*p*-values>0.224). (Individual tumor volumes are shown in Figures S16 and S17.)

**FIGURE 5.**
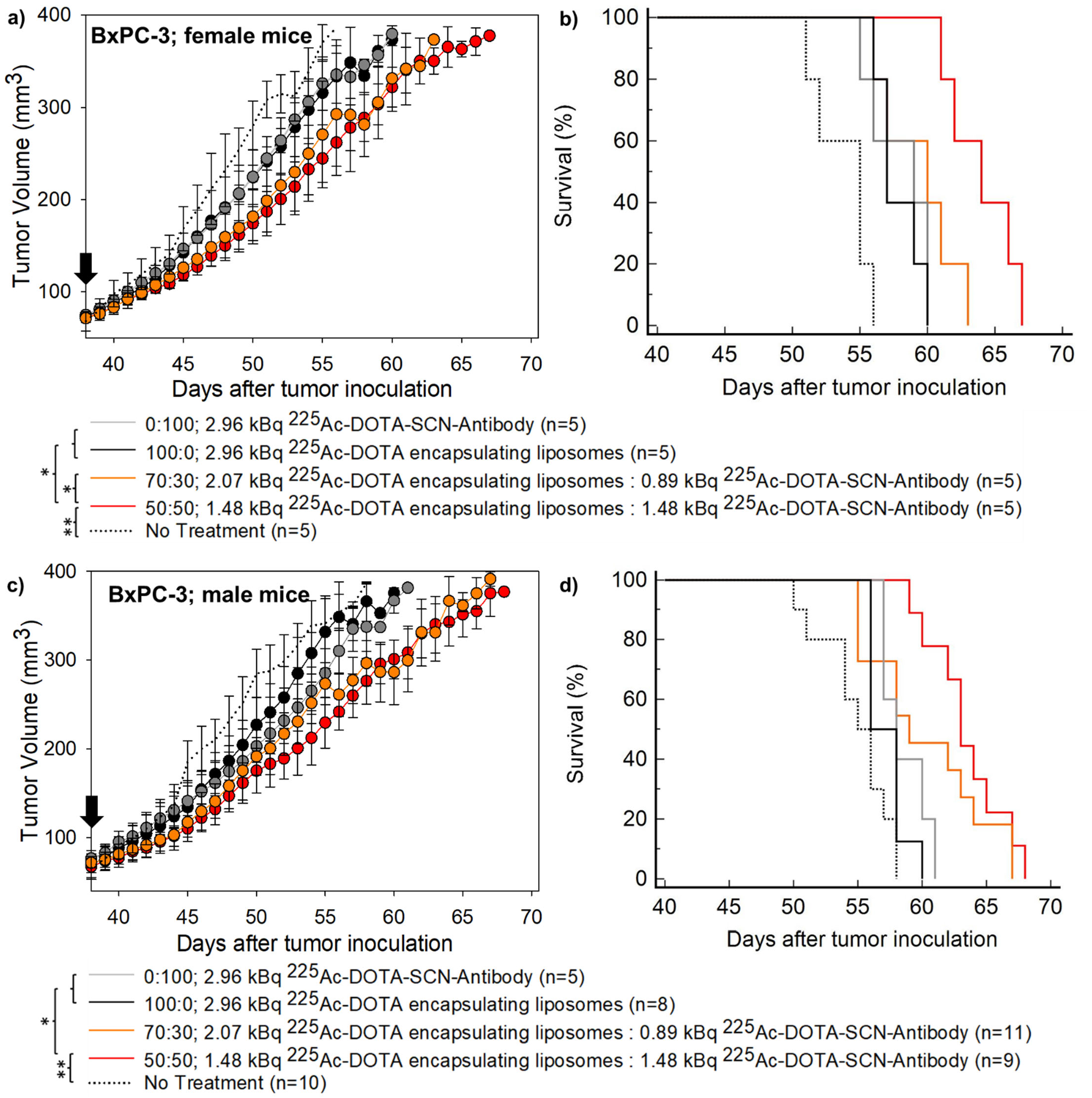
Growth inhibition of (too) low HER1-expressing, BxPC3 pancreatic tumors on female (a) and male (c) NSG mice bearing subcutaneous xenografts, and the corresponding survival (b, d), following treatment with a single i.v. injection of 2.96 kBq ^225^Ac per 20 g mouse (indicated by the black vertical arrows in (a) and (c)) delivered by: the HER1-targeting Cetuximab (grey symbols), the tumor-responsive liposomes (black symbols), and/or both, separate carriers at different split ratios (red, orange colored symbols) of same total injected radioactivity. Error bars correspond to the standard deviation of measurements averaged over n mice per condition as indicated. Significance in survival was calculated with one-way ANOVA (*p*-value<0.05). Characteristic H&E-stained sections of tumors and normal organs are shown in Figure S22.

**FIGURE 6.**
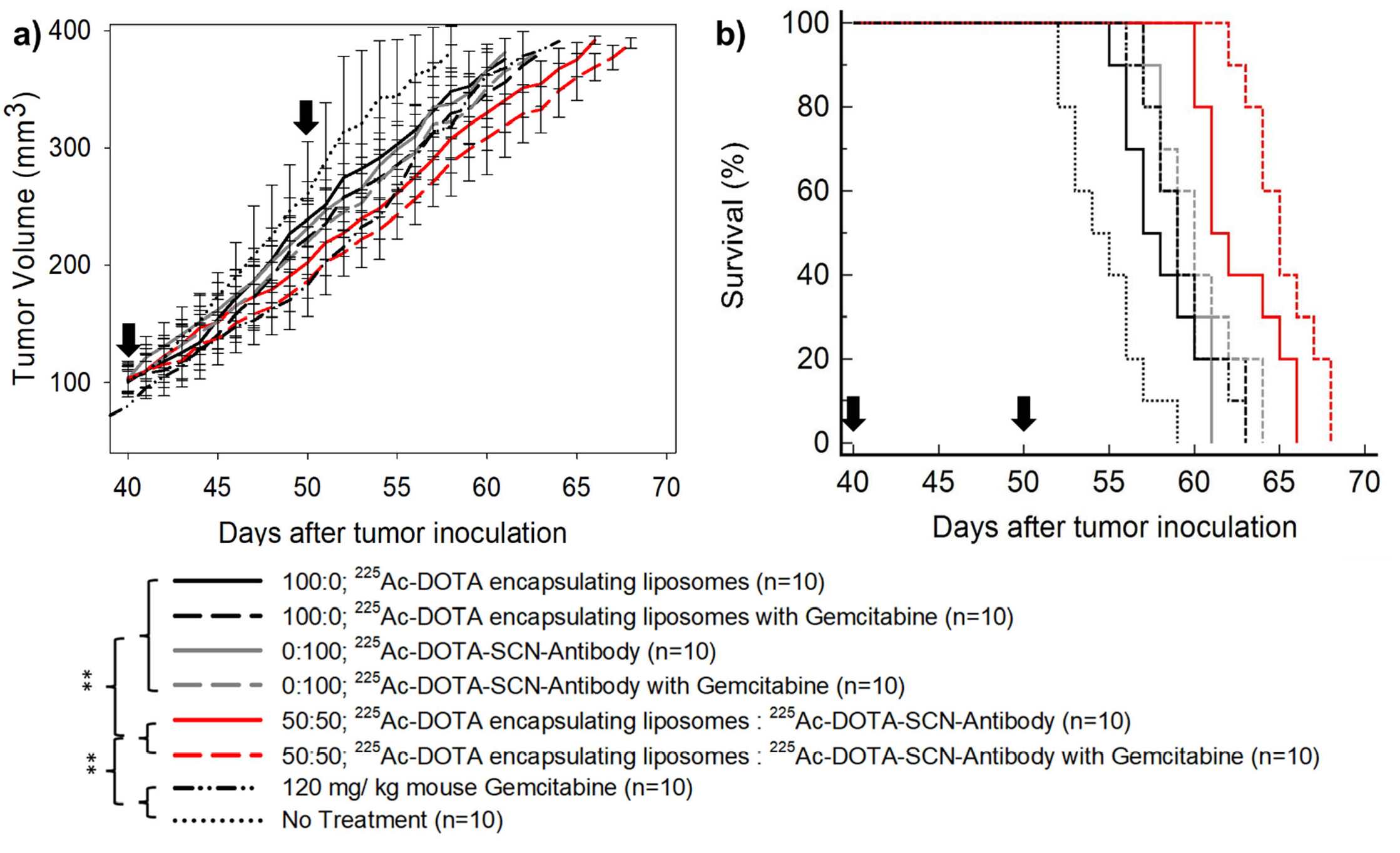
Inhibition of (a) tumor growth and (b) survival of NSG male mice bearing BxPC3 subcutaneous pancreatic tumors, following treatment with i.v. injected 2.22 kBq (on day 40 after tumor inoculation, black vertical arrows in (a) and (c)) and 0.74 kBq (on day 50) of ^225^Ac per 20 g male mouse delivered by: the HER1-targeting Cetuximab (grey lines), the tumor-responsive liposomes (black lines), and/or both, separate, carriers at 50:50 split ratio of radioactivity (red lines) in the absence (solid lines) and presence (dashed lines) of gemcitabine. Gemcitabine was injected intraperitoneally on day 38, after tumor inoculation, at 120 mg/Kg. Significance in survival was calculated with one-way ANOVA (*p*-value<0.05). Characteristic H&E-stained sections of tumors and normal organs are shown in Figure S23.

In both TNBC animal models, histopathology did not reveal off-target toxicities (Figures S18 and S19); only the spleen showed moderate to high hemosiderin deposition in the red pulp and reduced extramedullary hematopoiesis. Animal weight did not decrease beyond 10-12% of the original weight throughout the study (Figures S20 and S21). The endpoint of the experimental design for the two TNBC models forming spontaneous metastases was day 14-16 after injection of therapy. Our primary goal with breast cancer mouse models was to confirm the control of primary tumor growth and to detect the time of metastatic onset. The above endpoints were set by the longest possible period before ulceration of the primary tumor in untreated mice, necessitating euthanasia.

Similarly, in the pancreatic animal model, pathology evaluation did not reveal toxicity due to radiation in off-target organs (Figures S22 and S23), and animal weights for all cohorts were maintained within 10-15% of the original mouse weight (Figures S24 and S25).

## Discussion

The treatment of tumors with low expression of markers targeted with α-particle-emitting radionuclide-antibody conjugates is promising (*20*). However, even if the high killing potential of α-particles proves these treatment designs successful, the relatively low tumor marker expression is not expected to entirely bypass the antibodies’ binding site barrier effect (*8,11,21,22*), therefore, still limiting their infiltration within solid tumors, in addition to delivering lower tumor absorbed doses. The heterogeneity in antibody microdistributions was also shown herein on low targeted- marker expressing spheroids, representing the tumor avascular regions. For these internalizing antibodies, as is also the case for the Thorium-labelled Trastuzumab currently in clinical trials, the resulting nonuniform intratumoral microdistributions are expected to not allow α-particle therapy to reach its full potential (*1,8,19*).

Our two-carrier approach aimed to alleviate the heterogeneity in intratumoral radionuclide microdistributions by combining two diverse carriers while keeping the injected total radioactivity unchanged. Combined with our reports on solid tumors overexpressing the targeted marker(s) (*1,8*), the present study validates the two-carrier strategy as a tumor-agnostic therapeutic intervention even for chemoresistant soft-tissue solid tumors with a wide range of expression levels of the targeted markers by cancer cells (IHC score from 1+ to 3+).

Regarding the applicability of this approach, scaling up, from µm-long spheroids and mm-long tumors, in mice, to cm-long tumors in humans, is not expected to significantly alter the general trends in efficacy. This is because the intratumoral distributions of drugs are not determined so much by the overall size of tumors but, rather, by the tumor avascular distances (distances to the nearest blood vessels within the tumors). These distances are not very different between mice and humans, and range within the order of several hundredths of µm (*23*). However, the major limitations of the two-carrier approach are expected to be (1) the tumor vascular permeability to antibodies and tumor-responsive liposomes, which, in some cases, could be bypassed with additional interventions, such as ultrasound irradiation (*24*), and (2) the level of tumor acidification that affects the release property of liposomes. The property of content release by the nanoparticle carrier within the tumor interstitium is critical, as indicated in Figure S26. MRI-generated maps of tumor extracellular pH in a variety of mouse models revealed highly heterogeneous intratumoral acidity and variability among animals (*1,8,13*), which, however, did not seem to affect the efficacy of the tumor-responsive liposomes. In humans, tumor interstitial acidification is highly variable among different tumor types (*25*) and could be a factor limiting the efficacy of the present design of tumor-responsive liposomes. Notably, however, tumor-responsive liposomes are engineered to be triggered under slightly acidic conditions (pH ∼ 6.8-6.7), which enables relatively reasonable applicability of this design, given that the reported levels of acidification of humans’ tumor interstitia fall within the range of (pH_e_∼6.7-6.5 (*25*)). If the intratumoral pH gradient is inadequate to induce content release from liposomes (or from any other nanometer-sized carrier), heat- and/or light-activated release mechanisms may be employed as alternative triggers for release (*26*).

Regarding off-target toxicities, biodistribution studies confirmed the spleen to be the common critical normal organ for both carriers, and the heart for antibody delivery. No pathological concerns due to irradiation were observed, mostly because of the relatively low injected radioactivity that was adequate to elicit the observed responses. In the future, if greater injected doses are required, saturation of the spleen with cold liposomes might partly address their uptake. The heart uptake is specific to Cetuximab, and any other antibody can be chosen as the carrier of radioactivity targeted to measurable (but not necessarily high) levels of surface markers on cancer cells (*1,8*).

In the pancreatic cancer tumor model, gemcitabine was chosen because it serves as the cornerstone chemotherapy for this indication (*27*). Previous studies on α-particle therapy targeting the overexpressed HER1 surface marker in mice, combined with gemcitabine, reported improved survival (*28*). We did not observe an improvement in survival when we combined gemcitabine with the α-particle therapy delivered by each carrier alone, possibly dye to the low tumor delivered doses. Conversely, however, the two-carrier approach, possibly because of better intratumoral spreading of these low delivered doses, noticeably improved survival extending the applicability of combining chemo- with α-radiotherapy on tumors with moderate-to-low marker expression.

In summary, this study validated the importance of intratumoral spreading of delivered α-particle radiotherapy using a *flexible and simple two-carrier approach*, even in tumors with moderate to low expression levels of the targeted markers, for which α-particle radionuclide-antibody conjugates are currently being evaluated in the clinic.

## Conclusion

In conclusion, the findings of this study show that with better spreading of α-particle therapy within tumors, the impact of α-particles can be expanded to include treatment of solid tumors independent of classification limits set by cancer origin and/or marker expression.

## Acknowledgements

The authors thank Dr George Sgouros at Johns Hopkins University for help with the dosimetry calculations, and Ms Pooja Hariharan and Mr Rohit Chaudhari for assistance with animal handling. This work was partially supported by grants from the W.W. Smith Charitable Trust, the Allegheny Health Network-Johns Hopkins Cancer Research Fund, and the Elsa U. Pardee Foundation.

