## Supporting Information for "Combined, yet Separate: cocktails of carriers (not drugs) for α-particle therapy of solid tumors expressing moderate-to-low levels of targetable markers"

#### Materials and Methods

The lipids 1,2-diarachidoyl-sn-glycero-3-phosphocholine (20:0 PC), 1,2-dipalmitoyl-sn-glycero-3-phospho-L-serine (sodium salt) (DPPS), 1,2-dipalmitoyl-sn-glycero-3-phosphoethanolamine-N-(lissamine rhodamine B sulfonyl) (ammonium salt) (DPPE-rhodamine) and 1,2-distearoyl-sn-glycero-3-phosphoethanolamine-N-PEG2000-dimenthylammonium propane/propanoyl (DSPE-PEG(2000)-DAP) were purchased from Avanti Polar Lipids (Alabaster, AL, USA) and were used without further purification (>99% purity). Cholesterol, Phosphate Buffered Saline (PBS), Sephadex G-50, Sepharose-4B, chloroform, ascorbic acid, 4-(2-hydroxyethyl)-1-piperazineethanesulfonic acid (HEPES), sucrose, trisodium citrate dihydrate, anhydrous citric acid, poly(2-hydroxyethyl methacrylate) (polyHEMA), diethylenetriaminepentaacetic acid (DTPA), 1,4,7,10-tetraazacyclododecane-1,4,7,10-tetraacetic acid (DOTA) and the calcium ionophore A23187 were purchased from Sigma-Aldrich (Atlanta, GA, USA). Ethylenediaminetetraacetic acid (EDTA) was purchased from Fisher Scientific (Pittsburgh, PA, USA), trypsin and Matrigel™ were purchased from Corning (Corning, NY, USA), penicillin-streptomycin was from ThermoFisher Scientific (Waltham, MA, USA), S-2- (4-Isothiocyanatobenzyl)-diethylenetriamine pentaacetic acid (DTPA-SCN) and S-2-(4-isothiocyanatobenzyl)-1,4,7,10-tetraazacyclododecane-1,4,7,10-tetraacetic acid (DOTA-SCN) were from Macrocyclics (Dallas, TX, USA). Chelex® resin, chromatography and desalting columns from Bio-Rad (Hercules, CA, USA), syringe filters (0.22μm, Cat No. 76479-024) from VWR (Radnor, PA, USA), the polycarbonate membranes (100 nm) were purchased from GE Healthcare-Whatman (Chicago, IL, USA), the Dulbecco's Modified Eagle Medium (DMEM) and Roswell Park Memorial Institute (RPMI) medium were from ATCC (Manassas, VA, USA), the Fetal Bovine Serum (FBS) was from Omega Scientific (Tarzana, CA, USA), the 3-(4,5-dimethylthiazol-2-yl)-2,5-diphenyltetrazolium bromide (MTT) assay kit was purchased from Promega (Madison, WI, USA), and the HER1-binding antibody Cetuximab was purchased from Eli Lilly (Indianapolis, IN, USA). Indium-111 ($^{111}In$), indium chloride, was purchased from BWXT (Ontario, Canada).

*Spheroid spatiotemporal distributions and treatment*

The spatiotemporal profiles of the HER1-targeting antibody, the tumor-responsive liposomes (the carrier) and their, initially, encapsulated contents were evaluated by incubating 400 μm diameter spheroids with FITC-labeled HER1-targeting Cetuximab (0.06μM, ex/em: 494/518nm) or with tumor-responsive liposomes (2 mM lipid, containing 1 mole% DPPE-Rhodamine lipid, ex/em: 550/590 nm, and encapsulating 2 μM CFDA-SE, ex/em: 497/517nm, used as the drug surrogate). Spheroids were harvested at various timepoints - during incubation with the carriers (uptake) and upon being transferred in fresh media (clearance) -, were flash frozen in cryochrome, mounted on OCT gel and sectioned at 20 μm thickness. The equatorial slices were then imaged using a confocal fluorescence microscope (Zeiss LSM 780 Confocal Microscope, at 10X objective, White Plains, NY). To generate the corresponding calibration curves, known concentrations of Rhodamine-labeled lipid/liposomes, CFDA-SE and FITC-labeled antibody were measured, using the same microscope settings, in a quartz cuvette of 20 μm path-length. An in-house developed Matlab erosion code was applied on the images of spheroid sections to calculate the average intensity/concentration within each 5 μm-wide concentric ring of the section which was then plotted vs. its radial position, and the time-integrated, radial concentrations were calculated using the trapezoidal rule.

**Table S1**. Evaluation of the (a) the $K_D$ of the HER1-targeting antibody, Cetuximab, (b) the HER1-expression levels by cancer cells used in this study, and (c) the doubling time of each cell line used in the study. Shown are the mean values and standard deviations of repeated measurements.

| Cell line | PANC-1 | CR-MDA-MB-436 | MDA-MB-231 | BxPC-3 |
|---|---|---|---|---|
| **Measured $K_D$ (nM) ($^{111}$In-DTPA-SCN-Cetuximab)** (n=2) | 1.8 ± 0.2 | 0.5 ± 0.1 | 1.0 ± 0.2 | 0.3 ± 0.04 |
| **HER-1 receptor copies per cell** | 520,000 ± 12,000 | 258,000 ± 18,000 | 225,000 ± 21,000 | 117,000 ± 6,000 |
| **Doubling time (hours)** | 32 | 40 | 36 | 54 |

**Table S2**. Tabulated values of biodistributions of the HER1-targeting $^{111}$In-DTPA-SCN-antibody, Cetuximab, on all mouse models studied herein. Error bars correspond to standard deviations of n=3 mice per condition per time point.

1. Cetuximab BDs on the MDA-MB-436 orthotopic mouse model

| Time (hours) | **1** | **8** | **24** | **48** | **72** | **96** |
|---|---|---|---|---|---|---|
| Blood | 15.3 ± 3.67 | 11.76 ± 1.96 | 9.76 ± 2.76 | 7.05 ± 1.55 | 4.58 ± 0.38 | 3.09 ± 0.46 |
| Heart | 5.81 ± 0.25 | 3.77 ± 1.27 | 1.51 ± 0.28 | 1.28 ± 0.11 | 1.01 ± 0.12 | 0.74 ± 0.47 |
| Lungs | 0.06 ± 0.03 | 0.20 ± 0.09 | 0.13 ± 0.07 | 0.23 ± 0.10 | 0.24 ± 0.12 | 0.21 ± 0.08 |
| Liver | 8.60 ± 1.16 | 10.1 ± 1.98 | 10.2 ± 1.88 | 8.12 ± 2.05 | 4.35 ± 2.23 | 3.16 ± 1.03 |
| Spleen | 4.02 ± 1.23 | 8.66 ± 2.04 | 11.6 ± 0.53 | 10.3 ± 2.08 | 15.3 ± 1.48 | 8.11 ± 3.5 |
| Stomach | 1.12 ± 0.71 | 1.07 ± 0.44 | 0.36 ± 0.13 | 0.21 ± 0.09 | 0.15 ± 0.04 | 0.30 ± 0.20 |
| Intestines | 1.50 ± 1.18 | 5.61 ± 1.64 | 1.93 ± 1.07 | 1.21 ± 0.41 | 0.87 ± 0.46 | 0.29 ± 0.14 |
| Kidneys | 4.86 ± 1.01 | 3.99 ± 0.80 | 2.88 ± 1.25 | 1.18 ± 0.46 | 1.04 ± 0.43 | 0.07 ± 0.05 |
| Muscle | 0.18 ± 0.08 | 0.28 ± 0.07 | 0.15 ± 0.11 | 0.13 ± 0.06 | 0.08 ± 0.01 | 0.15 ± 0.07 |
| Bone | 0.15 ± 0.08 | 0.29 ± 0.15 | 0.12 ± 0.1 | 0.13 ± 0.06 | 0.05 ± 0.03 | 0.07 ± 0.04 |
| Tumor | 1.77 ± 1.56 | 3.09 ± 1.92 | 4.78 ± 0.98 | 5.45 ± 0.17 | 2.69 ± 0.56 | 2.21 ± 0.38 |

2. Cetuximab BDs on the MDA-MB-231 orthotopic mouse model

| Time (hours) | **1** | **8** | **24** | **48** | **72** | **96** |
|---|---|---|---|---|---|---|
| Blood | 15.9 ± 3.80 | 11.1 ± 0.54 | 8.91 ± 1.12 | 7.57 ± 1.28 | 4.32 ± 1.26 | 2.61 ± 0.64 |
| Heart | 5.80 ± 1.59 | 2.79 ± 1.43 | 1.74 ± 0.58 | 1.41 ± 0.29 | 1.46 ± 0.12 | 1.14 ± 0.24 |
| Lungs | 0.01 ± 0.00 | 0.84 ± 0.71 | 0.49 ± 0.41 | 0.99 ± 0.37 | 0.80 ± 0.47 | 0.32 ± 0.31 |
| Liver | 8.46 ± 1.89 | 11.70 ± 2.69 | 9.65 ± 0.96 | 8.76 ± 1.35 | 4.37 ± 2.26 | 2.98 ± 0.49 |
| Spleen | 4.76 ± 1.35 | 7.90 ± 0.50 | 10.20 ± 1.87 | 12.8 ± 0.98 | 15.90 ± 3.12 | 7.44 ± 2.13 |
| Stomach | 0.17 ± 0.01 | 0.86 ± 0.37 | 0.25 ± 0.20 | 0.25 ± 0.15 | 0.18 ± 0.11 | 0.02 ± 0.01 |
| Intestines | 2.28 ± 1.50 | 3.84 ± 0.97 | 2.09 ± 0.67 | 1.73 ± 0.24 | 1.13 ± 0.53 | 1.32 ± 0.07 |

| | | | | | | |
|---|---|---|---|---|---|---|
| Kidneys | 5.69 ± 1.13 | 3.27 ± 0.25 | 2.93 ± 0.62 | 1.35 ± 0.19 | 0.54 ± 0.25 | 0.36 ± 0.13 |
| Muscle | 0.08 ± 0.03 | 0.06 ± 0.02 | 0.03 ± 0.02 | 0.10 ± 0.06 | 0.08 ± 0.06 | 0.04 ± 0.03 |
| Bone | 0.02 ± 0.01 | 0.01 ± 0.01 | 0.01 ± 0.00 | 0.01 ± 0.01 | 0.07 ± 0.06 | 0.02 ± 0.01 |
| Tumor | 1.01 ± 0.61 | 2.81 ± 1.40 | 4.89 ± 1.29 | 6.19 ± 2.17 | 3.41 ± 1.75 | 2.64 ± 0.49 |

3. Cetuximab BDs on the BxPC-3 Female subcutaneous mouse model

| Time (hours) | **1** | **8** | **24** | **48** | **72** | **96** |
|---|---|---|---|---|---|---|
| Blood | 14.4 ± 3.71 | 11.20 ± 1.60 | 9.63 ± 0.21 | 6.50 ± 0.18 | 5.04 ± 0.17 | 3.28 ± 0.02 |
| Heart | 6.00 ± 0.81 | 3.23 ± 0.66 | 1.89 ± 0.23 | 1.35 ± 0.19 | 1.26 ± 0.16 | 1.10 ± 0.39 |
| Lungs | 0.02 ± 0.10 | 0.12 ± 0.02 | 0.19 ± 0.04 | 0.16 ± 0.03 | 0.13 ± 0.04 | 0.18 ± 0.03 |
| Liver | 6.57 ± 1.97 | 9.49 ± 1.27 | 14.40 ± 0.91 | 8.47 ± 1.06 | 4.81 ± 1.15 | 2.77 ± 0.36 |
| Spleen | 4.84 ± 1.03 | 7.34 ± 2.97 | 10.20 ± 0.63 | 10.3 ± 0.27 | 19.6 ± 3.23 | 7.44 ± 1.93 |
| Stomach | 1.20 ± 0.84 | 0.62 ± 0.25 | 0.39 ± 0.05 | 0.30 ± 0.03 | 0.21 ± 0.02 | 0.24 ± 0.03 |
| Intestines | 1.43 ± 0.80 | 6.44 ± 1.39 | 2.23 ± 1.18 | 1.52 ± 0.50 | 1.25 ± 0.63 | 0.46 ± 0.10 |
| Kidneys | 4.96 ± 1.58 | 2.29 ± 1.18 | 1.81 ± 0.67 | 1.27 ± 0.25 | 1.70 ± 0.17 | 0.05 ± 0.02 |
| Muscle | 0.10 ± 0.03 | 0.10 ± 0.04 | 0.14 ± 0.09 | 0.09 ± 0.03 | 0.09 ± 0.00 | 0.10 ± 0.01 |
| Bone | 0.08 ± 0.01 | 0.08 ± 0.01 | 0.12 ± 0.08 | 0.08 ± 0.02 | 0.08 ± 0.03 | 0.08 ± 0.00 |
| Tumor | 1.66 ± 1.16 | 2.28 ± 1.72 | 3.09 ± 1.31 | 3.29 ± 1.09 | 1.93 ± 0.96 | 1.65 ± 0.44 |

4. Cetuximab BDs on the BxPC-3 Male subcutaneous mouse model

| Time (hours) | **1** | **8** | **24** | **48** | **72** | **96** |
|---|---|---|---|---|---|---|
| Blood | 14.70 ± 1.12 | 11.70 ± 0.65 | 9.11 ± 1.49 | 7.23 ± 0.13 | 4.46 ± 0.69 | 2.69 ± 0.20 |
| Heart | 6.35 ± 1.93 | 3.36 ± 1.03 | 1.66 ± 0.33 | 1.30 ± 0.16 | 1.10 ± 0.60 | 0.65 ± 0.13 |
| Lungs | 0.09 ± 0.01 | 0.15 ± 0.03 | 0.18 ± 0.10 | 0.30 ± 0.05 | 0.26 ± 0.11 | 0.23 ± 0.09 |
| Liver | 7.47 ± 1.08 | 9.49 ± 1.24 | 12.3 ± 1.17 | 9.74 ± 2.18 | 4.70 ± 1.64 | 1.42 ± 0.49 |
| Spleen | 4.22 ± 0.73 | 8.91 ± 0.91 | 10.16 ± 1.87 | 13.60 ± 2.23 | 18.30 ± 2.58 | 9.03 ± 1.56 |
| Stomach | 1.41 ± 0.18 | 1.13 ± 0.41 | 0.67 ± 0.06 | 0.34 ± 0.04 | 0.29 ± 0.08 | 0.13 ± 0.09 |
| Intestines | 1.87 ± 0.73 | 3.38 ± 1.95 | 1.40 ± 0.13 | 1.28 ± 0.10 | 0.78 ± 0.20 | 0.16 ± 0.09 |

| | | | | | | |
|---|---|---|---|---|---|---|
| Kidneys | 4.56 ± 1.11 | 4.00 ± 0.40 | 2.54 ± 0.45 | 1.24 ± 0.37 | 1.32 ± 0.10 | 0.06 ± 0.02 |
| Muscle | 0.17 ± 0.10 | 0.22 ± 0.09 | 0.12 ± 0.08 | 0.18 ± 0.08 | 0.04 ± 0.01 | 0.09 ± 0.02 |
| Bone | 0.15 ± 0.05 | 0.21 ± 0.13 | 0.11 ± 0.09 | 0.19 ± 0.09 | 0.04 ± 0.01 | 0.08 ± 0.01 |
| Tumor | 1.75 ± 0.89 | 2.06 ± 1.55 | 2.79 ± 1.39 | 3.33 ± 1.72 | 1.67 ± 1.35 | 1.52 ± 1.10 |

**Table S3**. Tabulated values of biodistributions of the tumor-responsive liposomes loaded with $^{111}$In-DTPA on all mouse models studied herein. Error bars correspond to standard deviations of n=3 mice per condition per time point.

1. Liposome BDs on the MDA-MB-436 orthotopic mouse model

| %IA/g | | | | | | | |
|---|---|---|---|---|---|---|---|
| Time (hours) | **1** | **8** | **16** | **24** | **32** | **48** | **72** |
| Blood | 27.6 ± 3.26 | 11.10±2.56 | 7.88 ± 1.09 | 5.65 ± 1.32 | 3.62 ± 0.09 | 1.46 ± 0.40 | 0.08 ± 0.01 |
| Heart | 6.12 ± 2.28 | 2.96 ± 1.07 | 1.93 ± 1.29 | 1.48 ± 0.77 | 1.04 ± 0.42 | 0.77 ± 0.16 | 0.43 ± 0.42 |
| Lungs | 0.29 ± 0.18 | 0.99 ± 0.71 | 0.57 ± 0.18 | 0.44 ± 0.15 | 0.01 ± 0.00 | 0.56 ± 0.17 | 0.00 ± 0.00 |
| Liver | 5.37 ± 2.23 | 6.11 ± 1.20 | 6.23 ± 0.79 | 8.22 ± 2.02 | 11.2 ± 1.84 | 6.52 ± 1.89 | 3.08 ± 1.77 |
| Spleen | 0.12 ± 0.07 | 1.09 ± 0.24 | 2.10 ± 0.80 | 4.40 ± 1.30 | 9.16 ± 1.23 | 5.88 ± 0.32 | 4.02 ± 1.66 |
| Stomach | 0.14 ± 0.09 | 0.21 ± 0.13 | 0.29 ± 0.17 | 0.12 ± 0.03 | 0.17 ± 0.08 | 0.24 ± 0.12 | 0.16 ± 0.05 |
| Intestines | 0.45 ± 0.04 | 0.42 ± 0.37 | 0.43 ± 0.35 | 0.19 ± 0.09 | 0.67 ± 0.45 | 0.42 ± 0.38 | 0.45 ± 0.05 |
| Kidneys | 1.35 ± 0.77 | 1.51 ± 0.33 | 1.74 ± 0.27 | 1.59 ± 0.53 | 1.58 ± 0.69 | 1.28 ± 0.63 | 0.96 ± 0.43 |
| Muscle | 0.15 ± 0.06 | 0.14 ± 0.10 | 0.09 ± 0.14 | 0.17 ± 0.03 | 0.12 ± 0.14 | 0.06 ± 0.05 | 0.24 ± 0.15 |
| Bone | 0.19 ± 0.05 | 0.13 ± 0.11 | 0.22 ± 0.10 | 0.14 ± 0.12 | 0.23 ± 0.10 | 0.25 ± 0.17 | 0.35 ± 0.31 |
| Tumor | 1.88 ± 0.92 | 2.08 ± 0.88 | 3.36 ± 0.66 | 5.70 ± 1.74 | 3.74 ± 0.49 | 2.77 ± 0.13 | 1.76 ± 0.52 |

2. Liposome BDs on the MDA-MB-231 orthotopic mouse model

| %IA/g | | | | | | | |
|---|---|---|---|---|---|---|---|
| Time (hours) | **1** | **8** | **16** | **24** | **32** | **48** | **72** |
| Blood | 28.80±3.60 | 11.00±2.52 | 7.69 ± 0.32 | 5.14 ± 1.01 | 3.90 ± 1.27 | 1.63 ± 0.43 | 0.52 ± 0.09 |
| Heart | 6.20 ± 2.41 | 2.38 ± 1.18 | 1.86 ± 1.18 | 1.97 ± 0.76 | 1.59 ± 0.36 | 1.55 ± 0.59 | 0.83 ± 0.09 |
| Lungs | 0.84 ± 0.54 | 1.40 ± 1.14 | 0.75 ± 0.48 | 1.56 ± 0.82 | 0.87 ± 0.08 | 1.01 ± 0.55 | 0.81 ± 0.11 |
| Liver | 5.70 ± 2.91 | 5.68 ± 1.54 | 6.86 ± 1.17 | 7.90 ± 0.56 | 11.00±2.50 | 5.51 ± 1.44 | 3.58 ± 0.28 |
| Spleen | 0.06 ± 0.03 | 0.57 ± 0.52 | 1.87 ± 0.67 | 2.73 ± 1.88 | 8.92 ± 0.86 | 6.76 ± 0.37 | 3.74 ± 0.55 |
| Stomach | 0.18 ± 0.13 | 0.16 ± 0.06 | 0.32 ± 0.09 | 0.22 ± 0.05 | 0.79 ± 0.54 | 2.06 ± 0.32 | 0.51 ± 0.36 |

| | | | | | | | |
|---|---|---|---|---|---|---|---|
| Intestines | 0.24 ± 0.18 | 0.26 ± 0.14 | 1.17 ± 0.75 | 1.52 ± 0.33 | 1.06 ± 0.66 | 0.93 ± 0.65 | 0.27 ± 0.14 |
| Kidneys | 2.49 ± 1.49 | 3.22 ± 0.75 | 1.61 ± 0.19 | 1.54 ± 0.86 | 1.84 ± 0.47 | 1.44 ± 0.86 | 0.68 ± 0.13 |
| Muscle | 0.56 ± 0.46 | 0.33 ± 0.14 | 0.54 ± 0.16 | 0.59 ± 0.32 | 0.91 ± 0.82 | 0.44 ± 0.11 | 0.53 ± 0.27 |
| Bone | 0.86 ± 0.68 | 0.49 ± 0.08 | 0.65 ± 0.54 | 0.92 ± 0.37 | 0.82 ± 0.39 | 1.32 ± 0.63 | 0.50 ± 0.08 |
| Tumor | 1.95 ± 0.22 | 2.15 ± 0.72 | 2.67 ± 0.52 | 5.12 ± 0.63 | 3.13 ± 0.28 | 2.87 ± 0.28 | 1.65 ± 0.32 |

3. Liposome BDs on the BxPC-3 subcutaneous Female mouse model

| %IA/g | | | | | | | |
|---|---|---|---|---|---|---|---|
| Time (hours) | **1** | **8** | **16** | **24** | **32** | **48** | **72** |
| Blood | 23.9 ± 2.54 | 9.71 ± 1.81 | 5.26 ± 1.71 | 3.08 ± 0.23 | 1.88 ± 0.05 | 0.23 ± 0.13 | 0.19 ± 0.01 |
| Heart | 5.77 ± 1.55 | 2.55 ± 1.10 | 1.72 ± 0.99 | 1.28 ± 0.72 | 0.94 ± 0.11 | 0.37 ± 0.08 | 0.26 ± 0.11 |
| Lungs | 0.17 ± 0.11 | 0.54 ± 0.16 | 0.65 ± 0.05 | 0.23 ± 0.04 | 0.67 ± 0.03 | 0.23 ± 0.14 | 0.01 ± 0.01 |
| Liver | 5.99 ± 2.36 | 6.22 ± 1.31 | 6.41 ± 0.8 | 8.46 ± 2.00 | 11.0 ± 0.59 | 7.32 ± 1.46 | 3.15 ± 0.82 |
| Spleen | 0.16 ± 0.05 | 1.21 ± 0.11 | 2.52 ± 0.99 | 5.15 ± 0.85 | 10.5 ± 0.61 | 5.61 ± 0.37 | 4.23 ± 1.03 |
| Stomach | 0.13 ± 0.09 | 0.24 ± 0.06 | 0.26 ± 0.18 | 0.12 ± 0.05 | 0.18 ± 0.03 | 0.10 ± 0.01 | 0.12 ± 0.01 |
| Intestines | 0.47 ± 0.07 | 0.49 ± 0.17 | 0.42 ± 0.25 | 0.43 ± 0.12 | 0.60 ± 0.24 | 0.42 ± 0.17 | 0.30 ± 0.11 |
| Kidneys | 1.28 ± 0.64 | 1.67 ± 0.60 | 1.87 ± 0.40 | 1.30 ± 0.40 | 1.01 ± 0.41 | 0.76 ± 0.40 | 0.50 ± 0.22 |
| Muscle | 0.11 ± 0.04 | 0.13 ± 0.05 | 0.25 ± 0.01 | 0.24 ± 0.04 | 0.28 ± 0.11 | 0.04 ± 0.03 | 0.15 ± 0.08 |
| Bone | 0.20 ± 0.04 | 0.15 ± 0.06 | 0.23 ± 0.01 | 0.18 ± 0.05 | 0.20 ± 0.12 | 0.24 ± 0.03 | 0.26 ± 0.09 |
| Tumor | 1.78 ± 0.71 | 2.08 ± 0.46 | 3.23 ± 0.28 | 5.18 ± 0.51 | 3.55 ± 0.45 | 1.82 ± 0.43 | 1.15 ± 0.10 |

4. Liposome BDs on the BxPC-3 subcutaneous Male mouse model

| %IA/g | | | | | | | |
|---|---|---|---|---|---|---|---|
| Time (hours) | **1** | **8** | **16** | **24** | **32** | **48** | **72** |
| Blood | 31.30±3.59 | 12.90±2.94 | 8.94 ± 1.58 | 4.94 ± 1.84 | 3.46 ± 0.04 | 1.57 ± 0.07 | 0.15 ± 0.02 |
| Heart | 5.76 ± 2.91 | 2.52 ± 1.68 | 1.53 ± 0.30 | 1.07 ± 0.42 | 0.61 ± 0.04 | 0.42 ± 0.07 | 0.16 ± 0.09 |
| Lungs | 0.30 ± 0.14 | 0.76 ± 0.32 | 0.79 ± 0.14 | 0.21 ± 0.11 | 0.46 ± 0.05 | 0.60 ± 0.04 | 0.09 ± 0.01 |
| Liver | 6.03 ± 1.73 | 6.80 ± 1.00 | 7.17 ± 0.89 | 8.21 ± 2.47 | 13.4 ± 0.45 | 7.12 ± 2.82 | 4.04 ± 1.52 |

| | | | | | | | |
|---|---|---|---|---|---|---|---|
| Spleen | 0.29 ± 0.07 | 1.26 ± 0.15 | 2.16 ± 0.46 | 5.16 ± 0.13 | 9.85 ± 0.38 | 5.36 ± 0.43 | 4.09 ± 1.09 |
| Stomach | 0.14 ± 0.04 | 0.30 ± 0.22 | 0.24 ± 0.04 | 0.19 ± 0.13 | 0.18 ± 0.11 | 0.35 ± 0.15 | 0.15 ± 0.09 |
| Intestines | 0.47 ± 0.07 | 0.41 ± 0.28 | 0.60 ± 0.38 | 0.43 ± 0.16 | 0.32 ± 0.07 | 0.52 ± 0.15 | 0.49 ± 0.12 |
| Kidneys | 1.07 ± 0.29 | 1.41 ± 0.48 | 1.32 ± 0.27 | 1.92 ± 0.31 | 1.85 ± 0.51 | 1.26 ± 0.18 | 0.83 ± 0.30 |
| Muscle | 0.15 ± 0.07 | 0.19 ± 0.09 | 0.17 ± 0.03 | 0.12 ± 0.09 | 0.26 ± 0.02 | 0.17 ± 0.08 | 0.15 ± 0.09 |
| Bone | 0.17 ± 0.08 | 0.18 ± 0.09 | 0.16 ± 0.03 | 0.12 ± 0.09 | 0.26 ± 0.02 | 0.17 ± 0.08 | 0.14 ± 0.09 |
| Tumor | 1.40 ± 0.17 | 1.73 ± 0.84 | 3.52 ± 0.89 | 5.34 ± 0.47 | 4.05 ± 0.39 | 1.94 ± 0.22 | 1.17 ± 0.10 |

**Figure S1**. Binding isotherms of the HER1-targeting $^{111}$In-DTPA-SCN-Cetuximab with each of the cancer cell lines used in this study. The immunoreactivity of the targeting antibody is indicated on each plot. The plotted concentration of the targeting antibody (on the x-axis) was corrected by multiplying the antibody concentration by its corresponding immunoreactivity. The y-axis was corrected for the non-specific binding of antibody (1, 2).

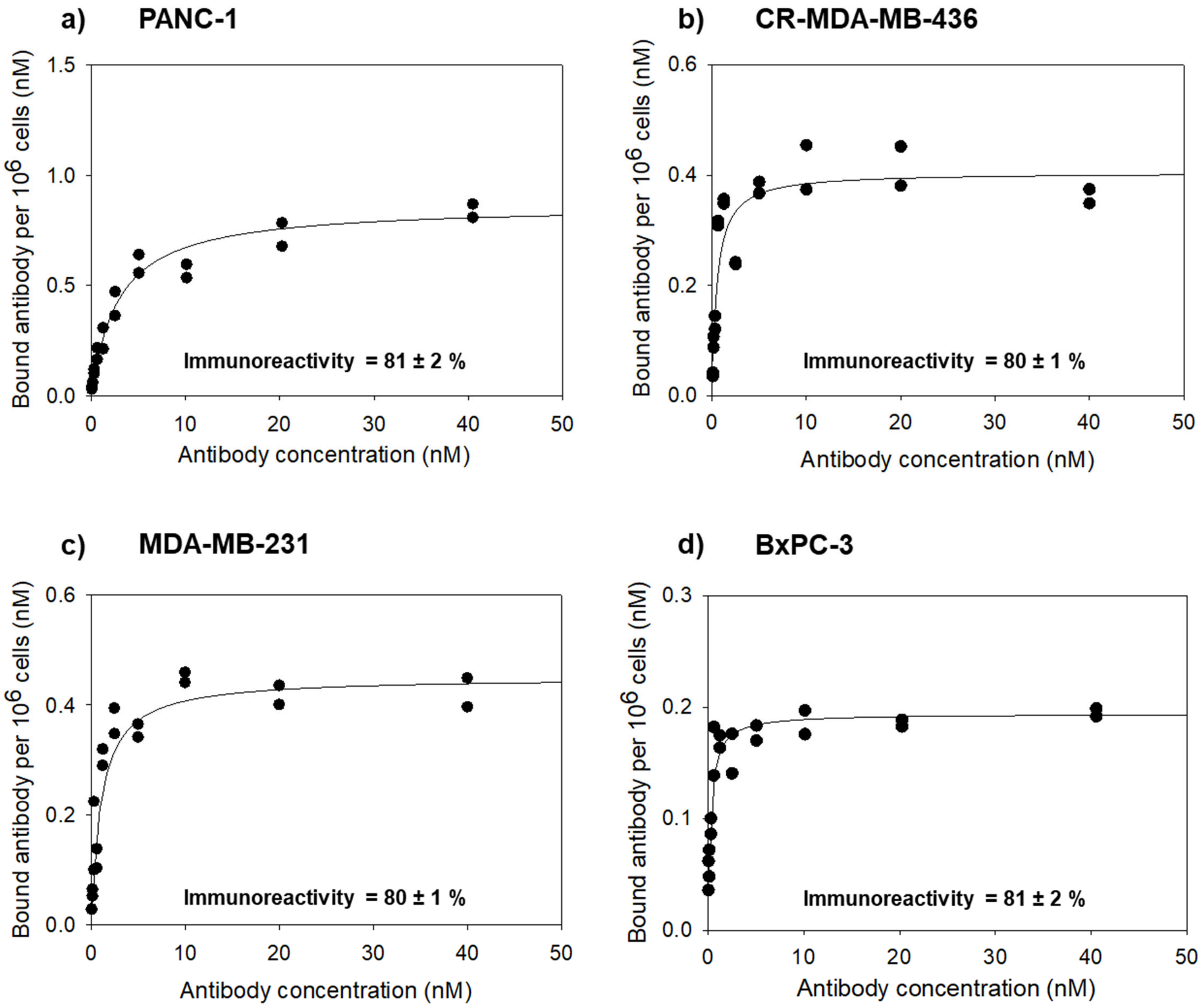

**Figure S2**. Flow cytometry of FITC-labelled Cetuximab incubated with the cancer cell lines studied herein. Red: cells only. Blue: cells incubated with fluorescent antibody.

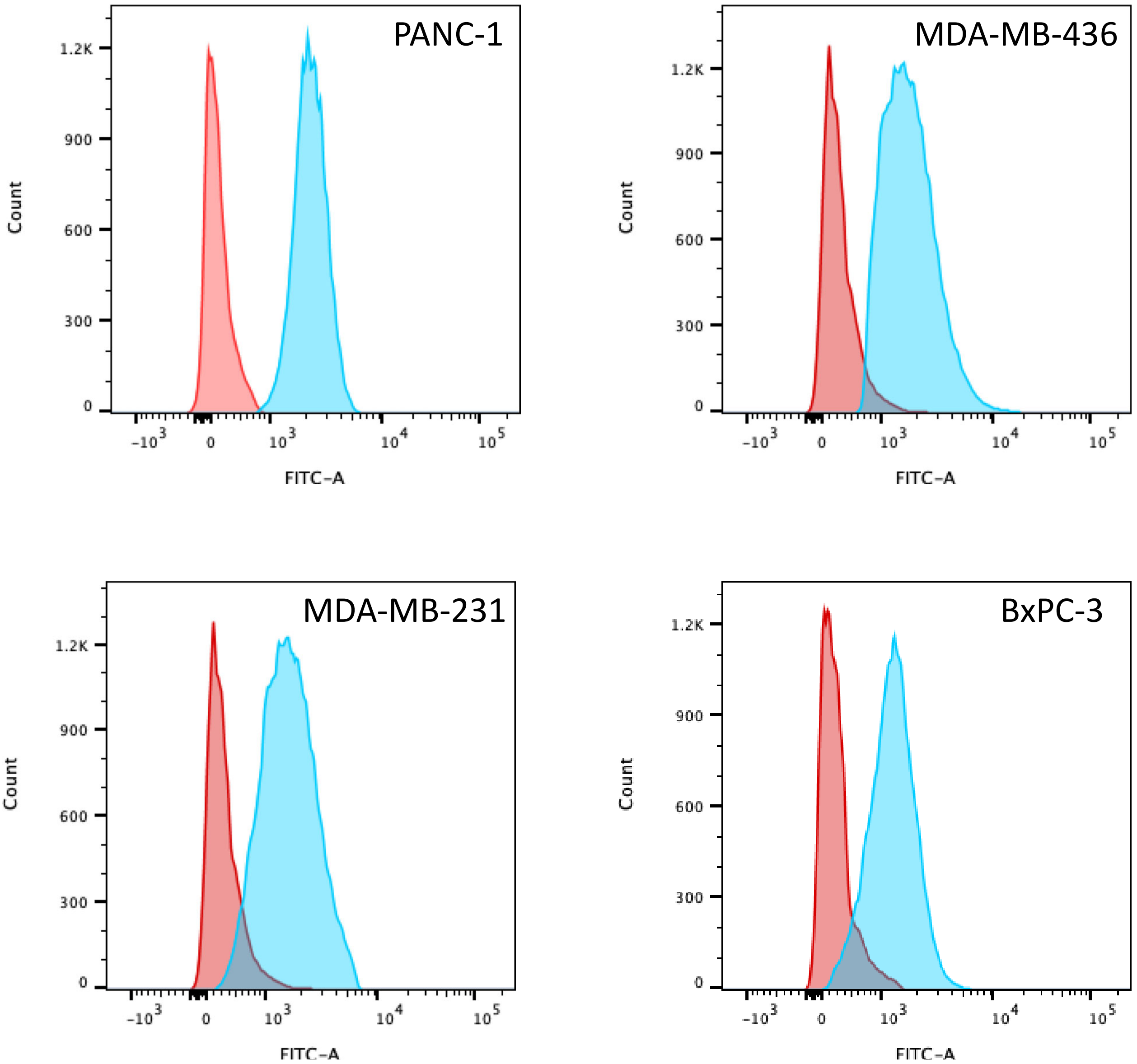


Method:

The cells were trypsinized and a suspension of 1 million cells/mL in media was obtained. For the fluorescent antibody conditions, this suspension was incubated with FITC-labeled Cetuximab at 50 times excess antibody-to-receptor ratio assuming 1 million receptors/cell on ice for 1 hour. Following this, the cell suspensions were centrifuged, the supernatant was removed, and the pelleted cells were resuspended in ice cold PBS. This procedure was repeated three times following which the cells were suspended in 1mL of PBS and were analyzed on the BD FACS Canto Flow cytometer (Franklin Lakes, NJ, USA). For the cells only condition, 1 million cells/mL were incubated with a volume of ice-cold PBS equal to the volume of fluorescently labeled antibody that was added.

**Figure S3**. Binding and internalization kinetics of the (fluorescently labeled) HER1-targeting antibody (Cetuximab) using each of the cell lines studied herein.

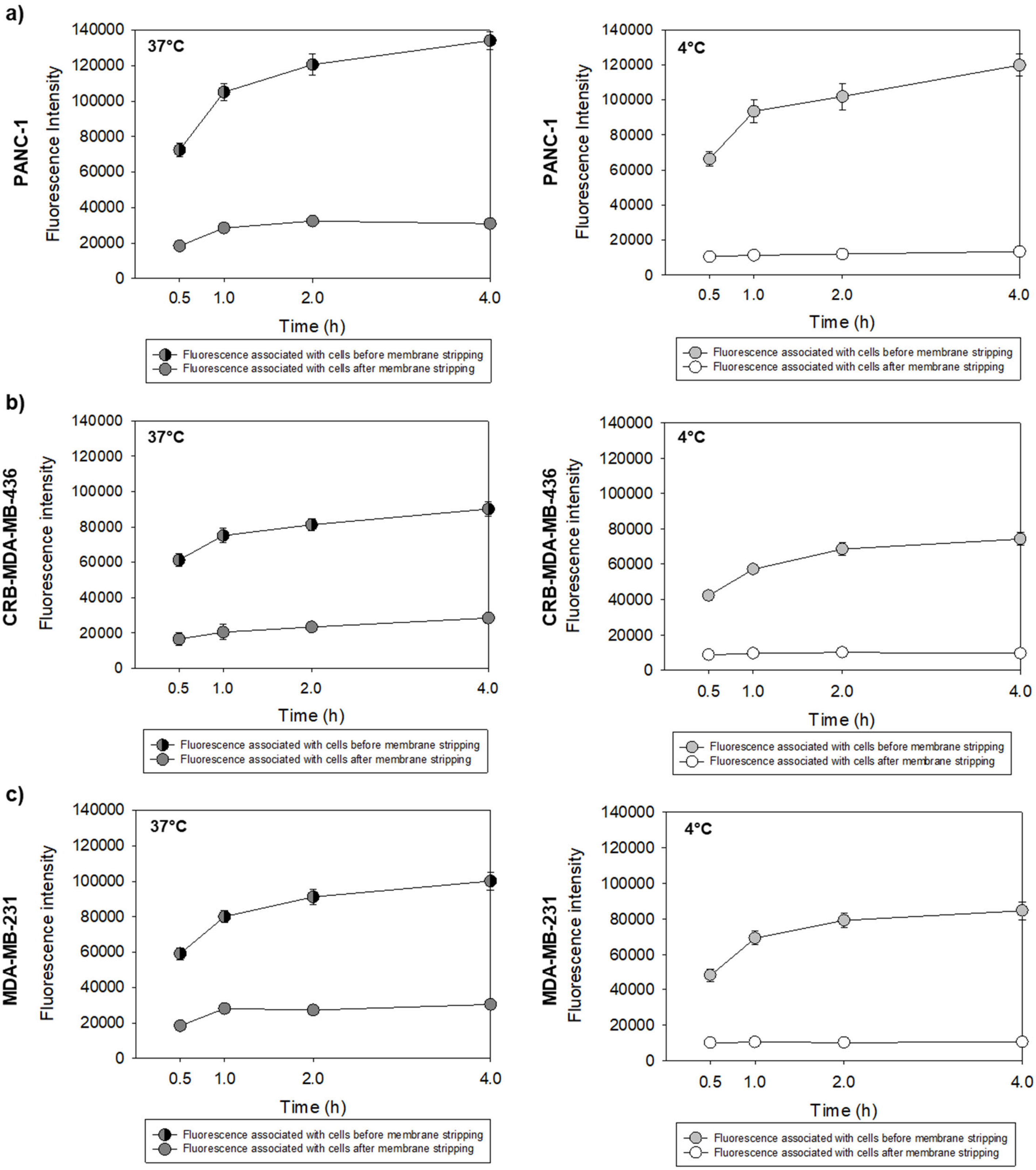

Method:

Cells were trypsinized and counted and were incubated with FITC-labeled HER1-targeting antibody (Cetuximab) at either 37 ºC or 4 ºC at a concentration of 2 million cells/mL. At 0.5, 1, 2 and 4 hours of incubation time, 1 mL of the parent suspension was removed, centrifuged and washed thrice with ice cold PBS. This suspension was mixed well and split into two halves of 0.5 mL each. For one half of the cell suspension, the fluorescence intensity associated with the cells (surface bound and internalized) was then measured, before and after cell lysis, to correct for any scattering effects from cells. The second part of the cell suspension was washed and incubated with acidic glycine buffer (50 mM glycine/150 mM NaCl, pH 3.0) for 10 minutes at room temperature to strip away the surface-bound antibody. The fluorescence intensity of this fraction (internalized antibody) was then measured as above. All counts were corrected for the number of cells and background intensity of cells.

**Figure S4**. Dose response curves of wild type MDA-MB-436 cells and their Cisplatin Resistant CR-MDA-MB-436 counterparts, on exposure to increasing concentrations of cisplatin at pH 7.4 (a) and pH 6.0 (b). $IC_{50}$ concentration of the resistant cells (white crossed circles) was evaluated to be almost 10-fold of that of the parental cells (black circles) at each pH (*, ǂ *p*-value<0.05). Symbols correspond to the mean values, and errors to the standard deviations of n=3 independent measurements.

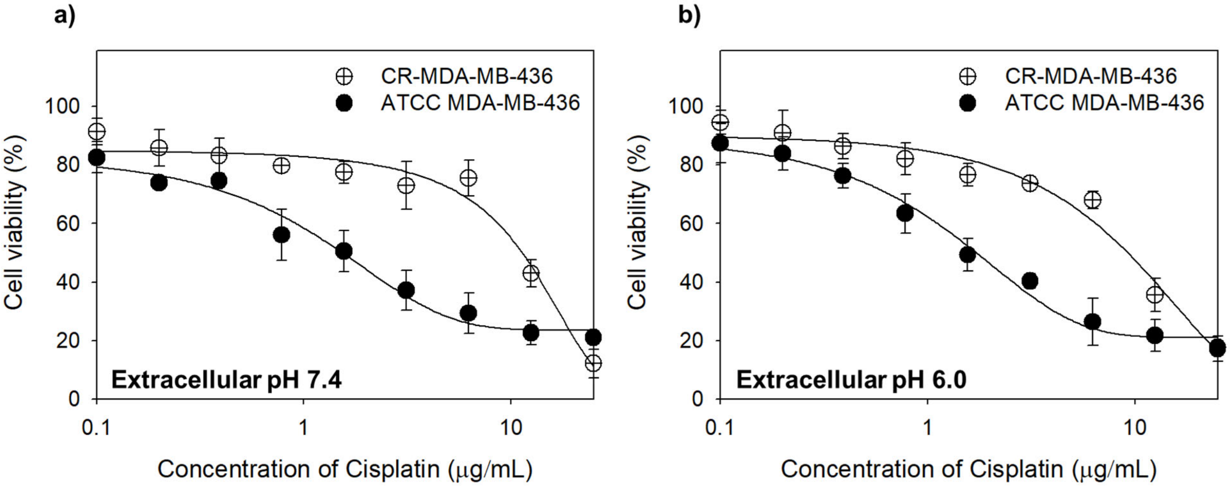


| Cell Line | Cisplatin $IC_{50}$ value (μg/mL) | |
|---|---|---|
| | pH 7.4 | pH 6.0 |
| Wild type MDA-MB-436 | 1.4 ± 0.2* | 1.2 ± 0.3ǂ |
| CR-MDA-MB-436 | 10.2 ± 2.9* | 8.2 ± 1.8ǂ |

Method: The Cisplatin $IC_{50}$ value (i.e., the concentration of Cisplatin at which 50% of the cells survive) was obtained by plating 20,000 cells/well the previous night on a 96-well flat-bottom plate. Cells were incubated with the Cisplatin-containing media (RPMI media that was preincubated in a humidified incubator at 37°C and 5% $CO_2$ overnight at pH 7.4 or 6.5 to ensure it was fully equilibrated) at different concentrations for six hours (3 wells per concentration at each pH). Following this treatment, cells were washed thrice with sterile PBS and then allowed to grow with fresh media for ~3.5 days (corresponding to two doubling times, 80 hours). On completion of two doubling times, the MTT assay was used to evaluate cell viability.

**Figure S5**. Colony survival of cancer cells following a 6-hour incubation at **pH=6.5** with different radioactivity concentrations of $^{225}$Ac-DOTA in free form (gray circles), $^{225}$Ac-DOTA delivered by tumor-responsive liposomes (black circles) and the HER1-targeting $^{225}$Ac-DOTA-SCN-Cetuximab (white circles). Colony survival response of cisplatin resistant CR-MDA-MB-436 cells (white crossed circles) was identical to their wild type counterparts (c), when exposed to $^{225}$Ac-DOTA. Data points indicate the mean values, and error bars the standard deviations of n = 3 independent cell-radioactivity incubation experiments.

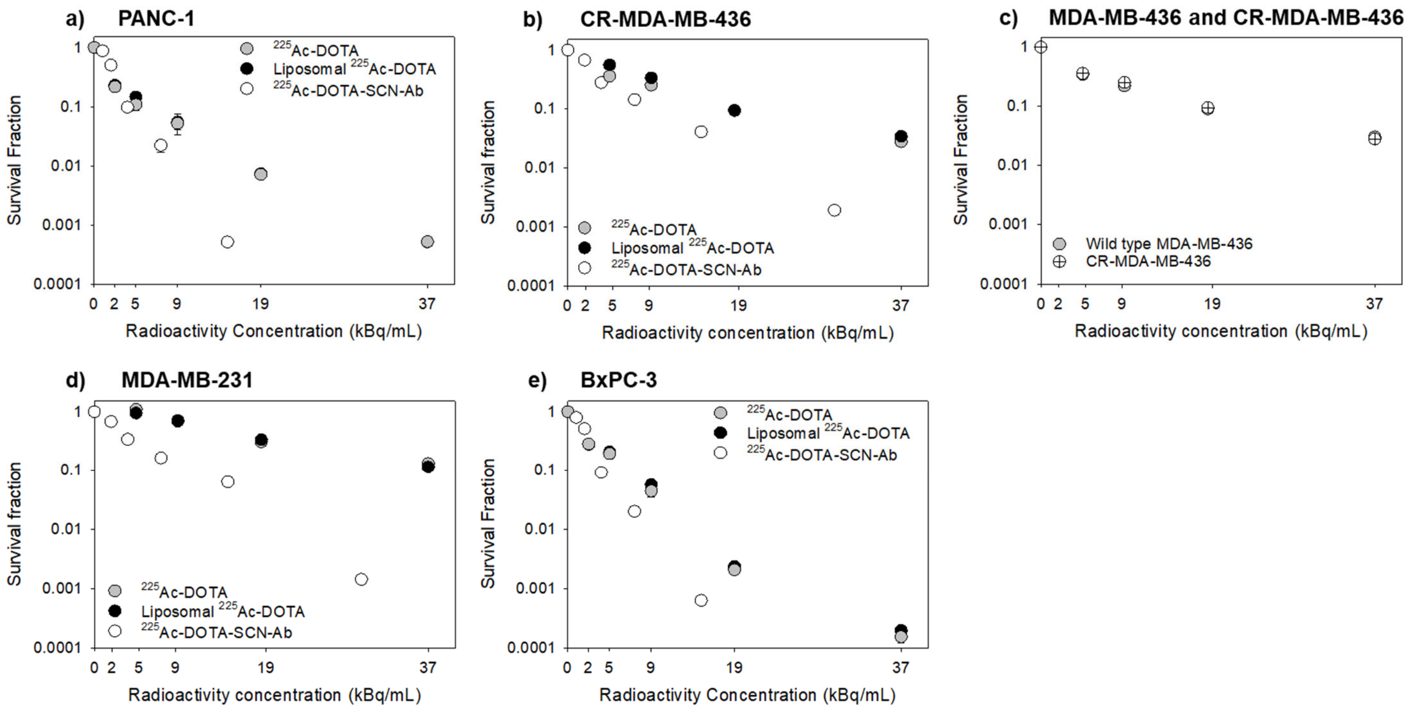

**Figure S6**. **Time-integrated spatiotemporal distributions** of the HER1-targeting antibody (a, d), the tumor-responsive liposomes (b, e), and of CFDA-SE, the fluorescent surrogate of $^{225}$Ac-DOTA delivered by liposomes (c, f), in spheroids formed by PANC-1 (top panel, used because it is a pancreatic cancer cell line that formed spheroids since BxPC3 did not form spheroids) and MDA-MB-231 (bottom panel) cells.

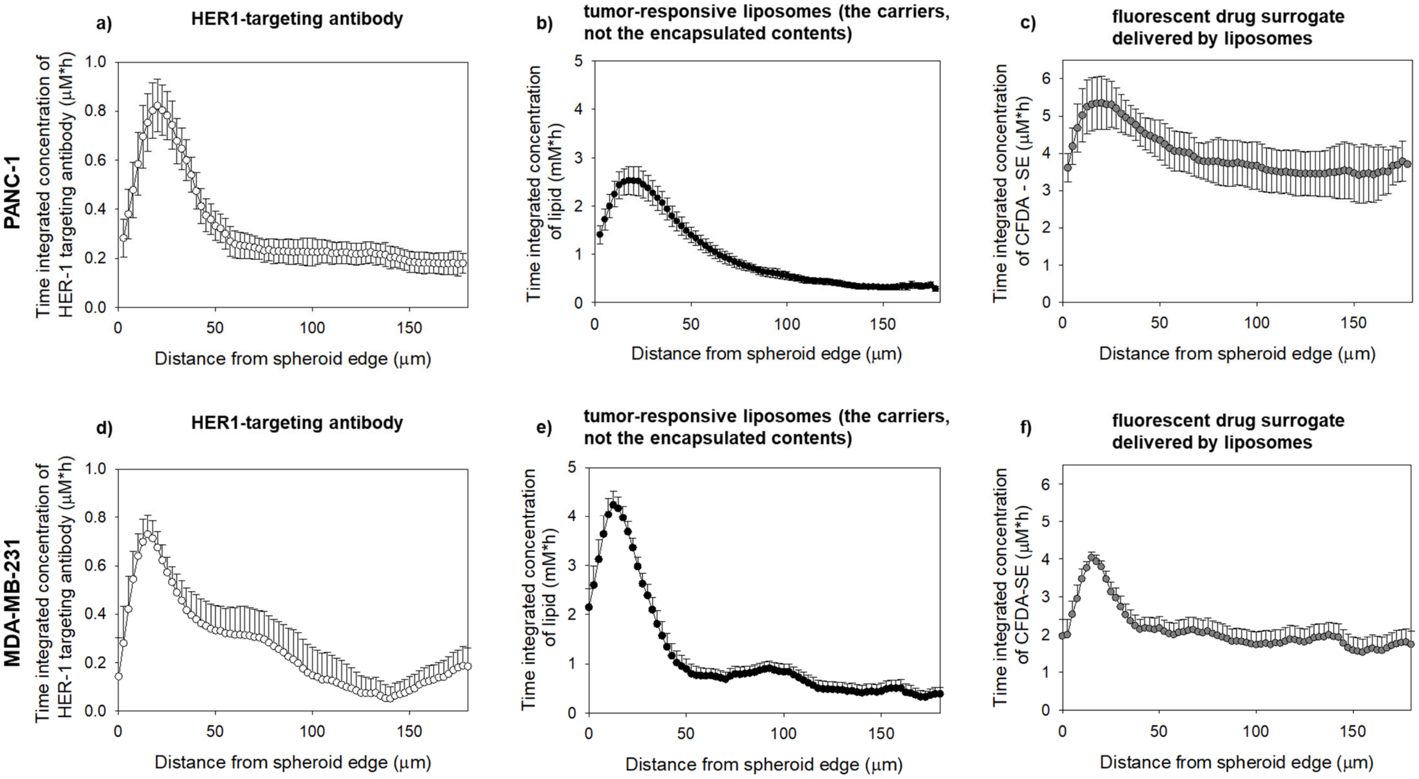

**Figure S7**. **Spatiotemporal profiles** of the HER1-targeting antibody (left column), Cetuximab; the tumor-responsive liposomes (middle); and of CFDA-SE (right column), the fluorescent surrogate of $^{225}$Ac-DOTA, delivered by liposomes in spheroids formed by **(a) PANC-1**, **(b) CR-MDA-MB-436**, and **(c) MDA-MB-231** cells. Liposomes contained fluorescently labeled lipids. During the "uptake" phase (upper raw), the corresponding carrier/drug surrogate was present in the surrounding medium at the following concentrations: 0.06 μM Cetuximab, 2 mM total lipid, 0.8 μM CFDA-SE (encapsulated in the tumor-responsive liposomes); during the "clearance" phase (lower raw), spheroids were transferred to media without any of the carriers/drug surrogate.

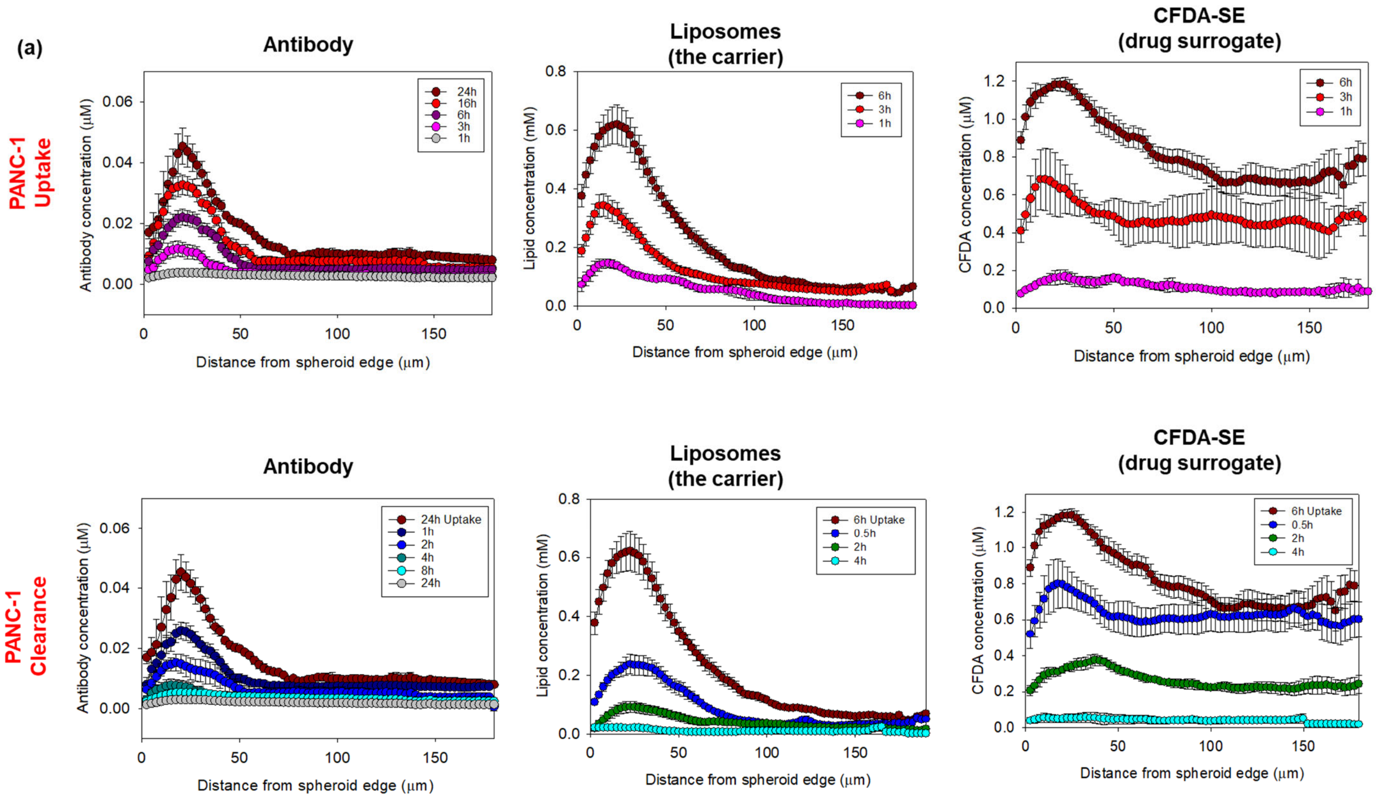

(b)

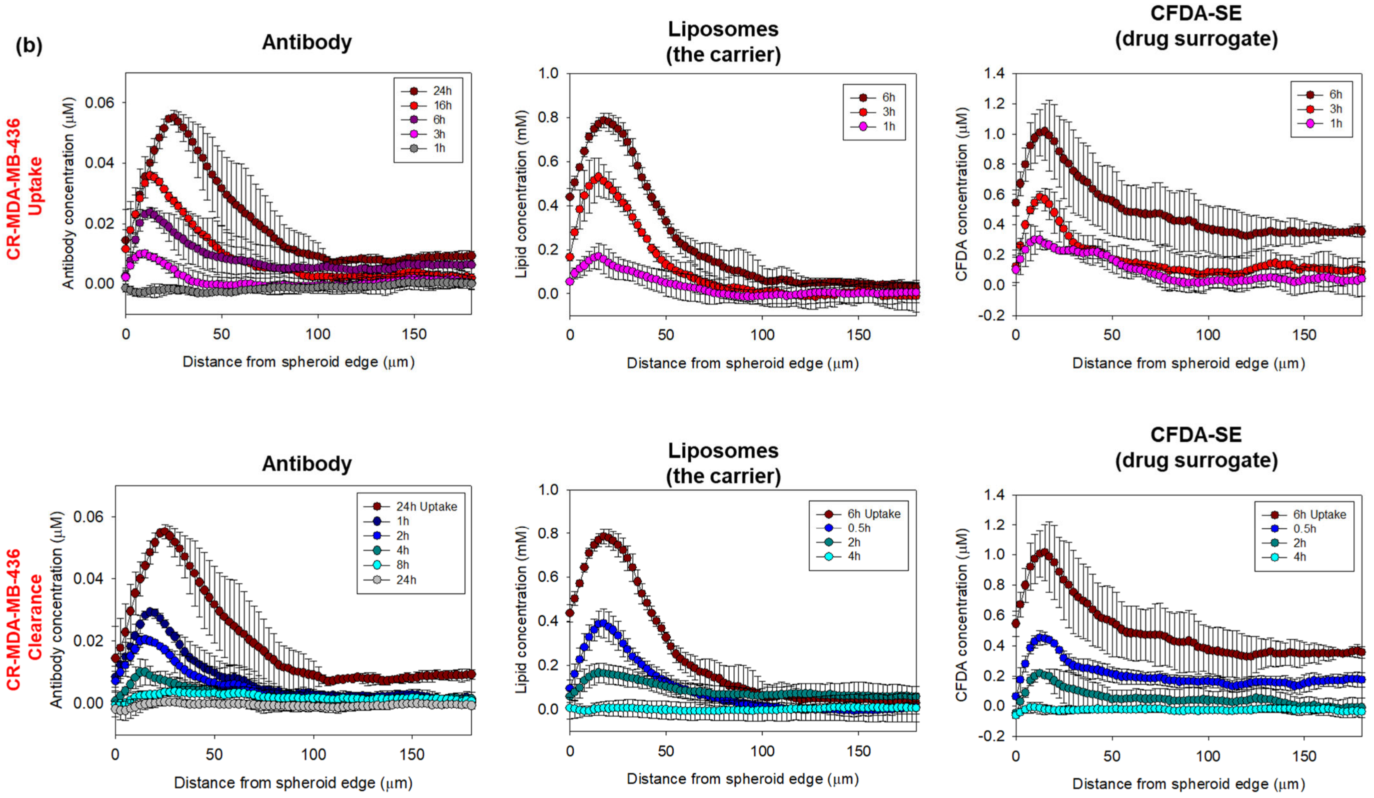

CR-MDA-MB-436 Uptake
Antibody
Antibody concentration (µM)
24h
16h
6h
3h
1h
Liposomes (the carrier)
Lipid concentration (mM)
6h
3h
1h
CFDA-SE (drug surrogate)
CFDA concentration (µM)
6h
3h
1h
CR-MDA-MB-436 Clearance
Antibody
Antibody concentration (µM)
24h Uptake
1h
2h
4h
8h
24h
Liposomes (the carrier)
Lipid concentration (mM)
6h Uptake
0.5h
2h
4h
CFDA-SE (drug surrogate)
CFDA concentration (µM)
6h Uptake
0.5h
2h
4h
Distance from spheroid edge (µm)


(c)

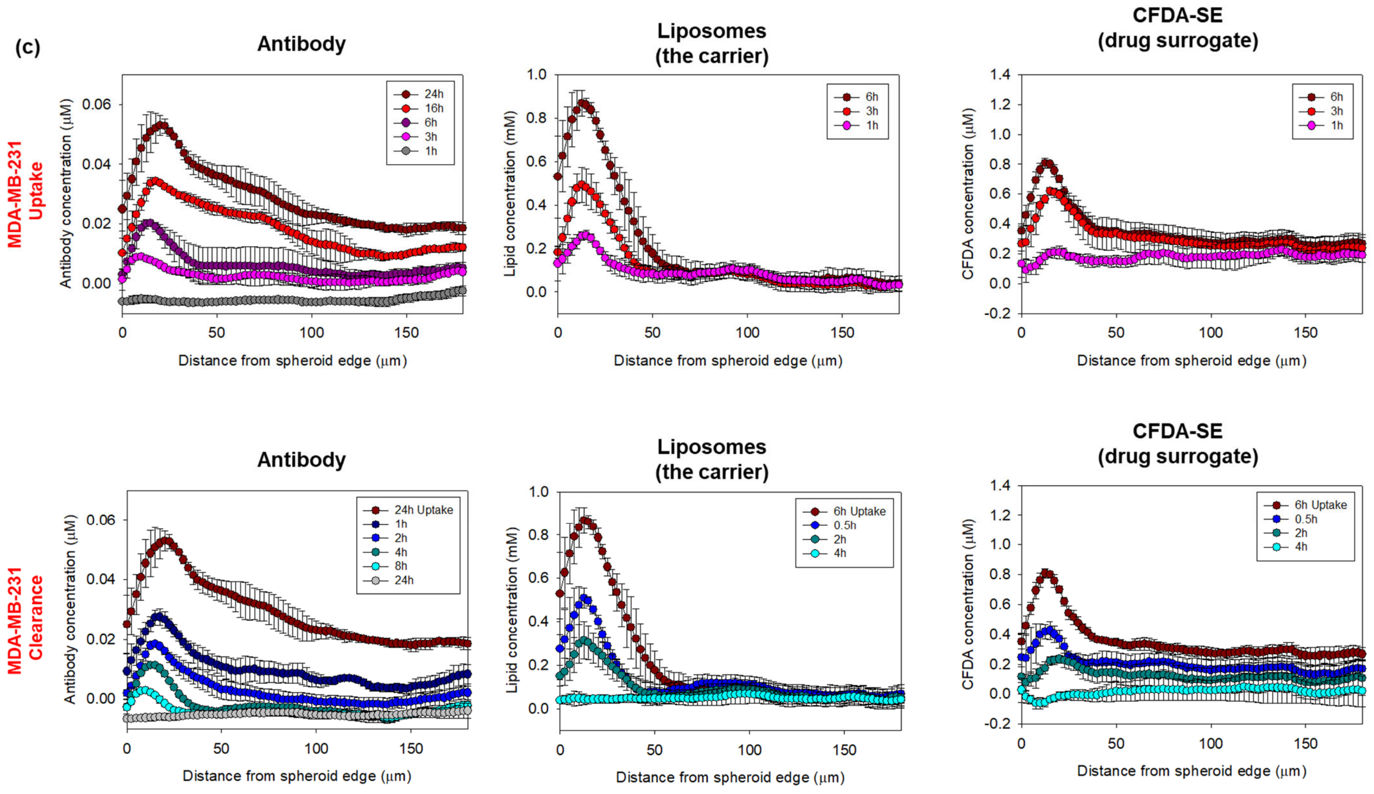

MDA-MB-231 Uptake
Antibody
Antibody concentration (µM)
24h
16h
6h
3h
1h
Liposomes (the carrier)
Lipid concentration (mM)
6h
3h
1h
CFDA-SE (drug surrogate)
CFDA concentration (µM)
6h
3h
1h
MDA-MB-231 Clearance
Antibody
Antibody concentration (µM)
24h Uptake
1h
2h
4h
8h
24h
Liposomes (the carrier)
Lipid concentration (mM)
6h Uptake
0.5h
2h
4h
CFDA-SE (drug surrogate)
CFDA concentration (µM)
6h Uptake
0.5h
2h
4h
Distance from spheroid edge (µm)

**Figure S8**. **Extent of outgrowth/regrowth inhibition** (used as indirect surrogate of tumor recurrence) of PANC-1 (top panel) and of MDA-MB-231 (bottom panel) spheroids of varying sizes at the time of treatment. The total radioactivity concentration was kept constant at 3.7 kBq/mL, and was divided at different ratios between the tumor-responsive liposomes loaded with $^{225}$Ac-DOTA (incubated for 6 hours) and the HER1-targeting $^{225}$Ac-DOTA-SCN-antibody (incubated for 24 hours). Error bars correspond to standard deviations of repeated measurements (n=12 spheroids per condition, n=2 independent liposome and antibody preparations). * *p*-values < 0.05.

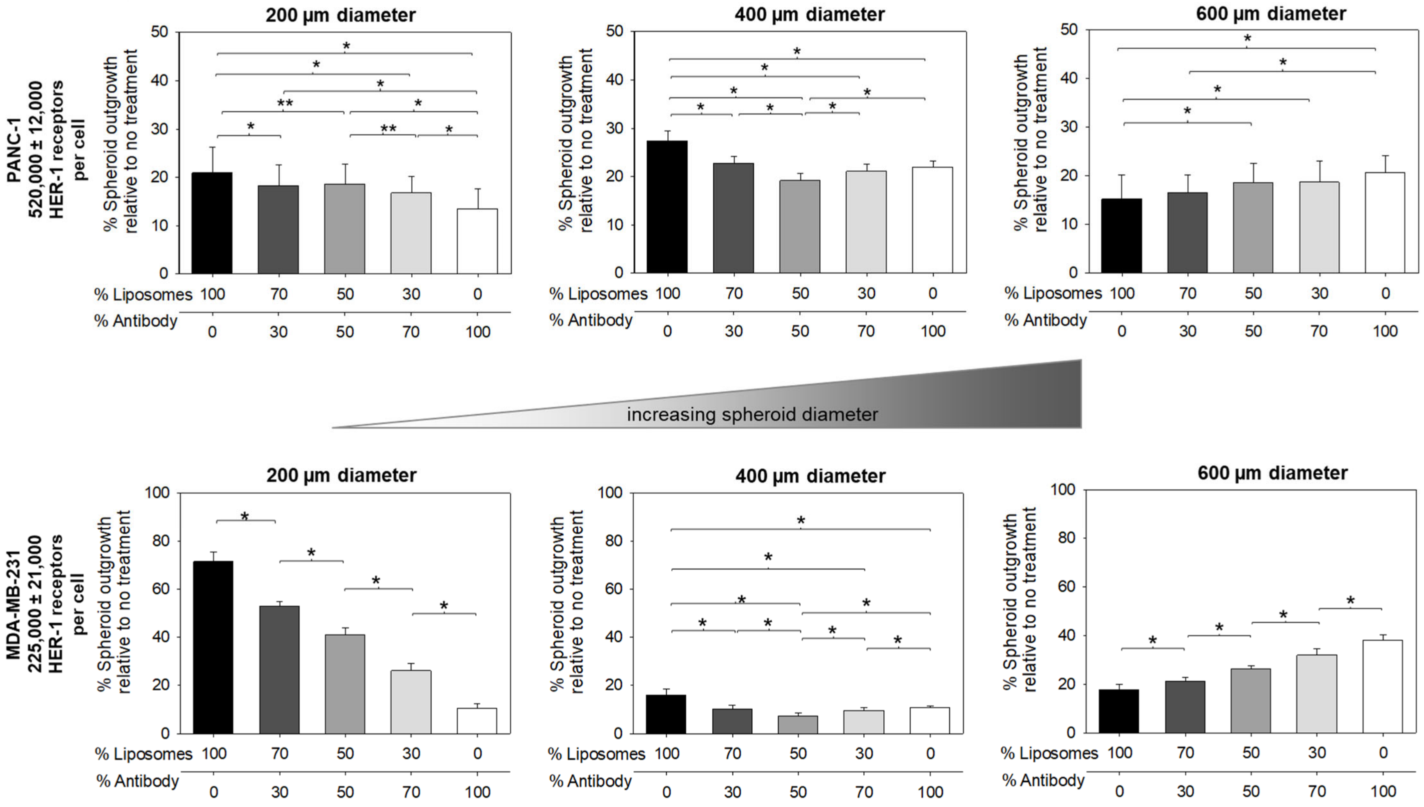

**Figure S9**. Biodistributions of (a) the HER1-targeting $^{111}$In-DTPA-SCN-antibody, Cetuximab, and the (b) tumor-responsive liposomes loaded with $^{111}$In-DTPA on NSG mice bearing **CR-MDA-MB-436** tumors. $^{111}$In was used as surrogate of the parent $^{225}$Ac. Error bars correspond to standard deviations of n=3 mice per condition per time point.

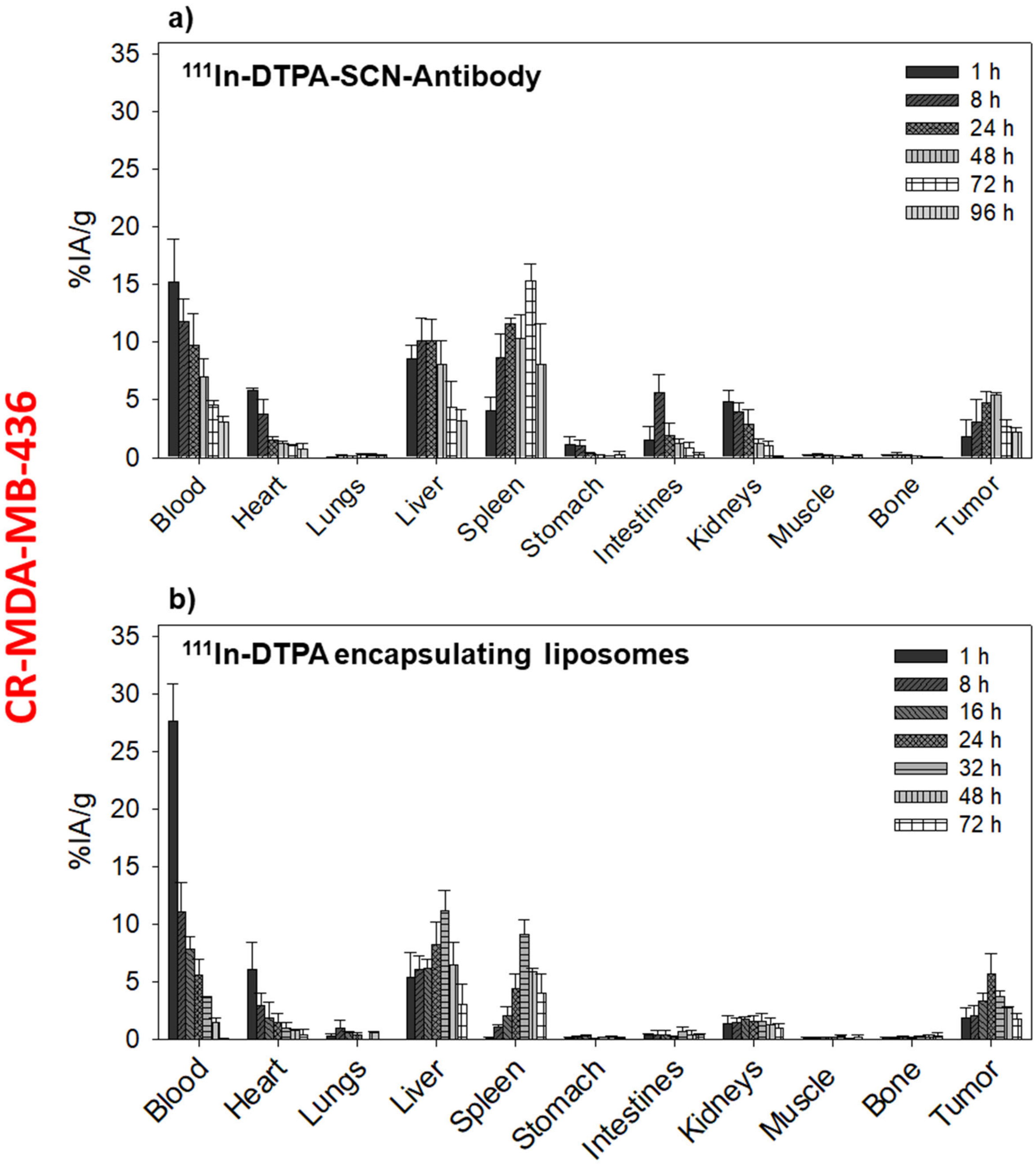

**Figure S10**. Biodistributions of (a) the HER1-targeting $^{111}$In-DTPA-SCN-antibody, Cetuximab, and the (b) tumor-responsive liposomes loaded with $^{111}$In-DTPA on NSG mice bearing **MDA-MB-231** tumors. $^{111}$In was used as surrogate of the parent $^{225}$Ac. Error bars correspond to standard deviations of n=3 mice per condition per time point.

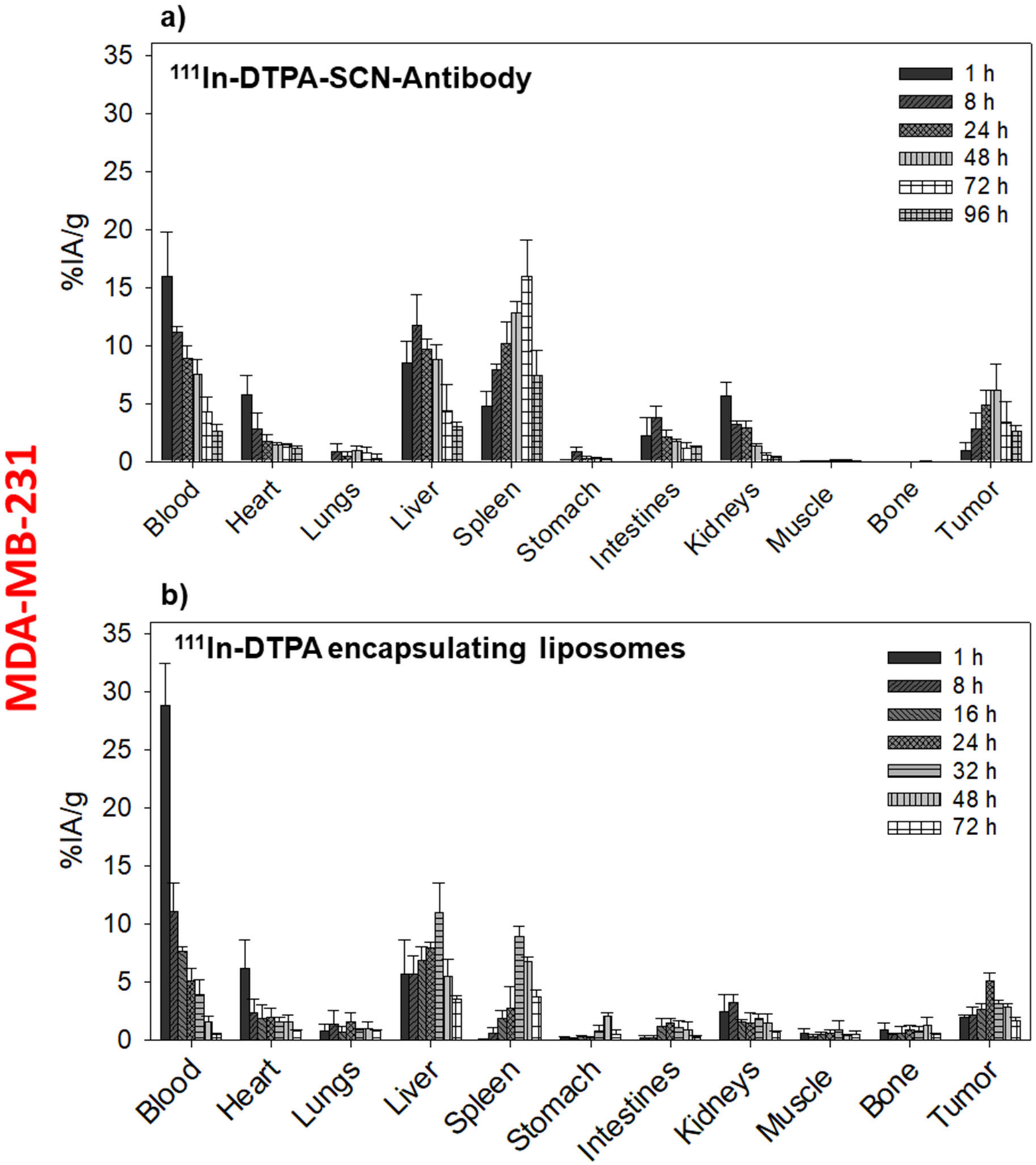

**Figure S11**. Biodistributions of (a) the HER1-targeting $^{111}$In-DTPA-SCN-antibody, Cetuximab, and the (b) tumor-responsive liposomes loaded with $^{111}$In-DTPA on **NSG female mice bearing BxPC3** tumors. $^{111}$In was used as surrogate of the parent $^{225}$Ac. Error bars correspond to standard deviations of n=3 mice per condition per time point.

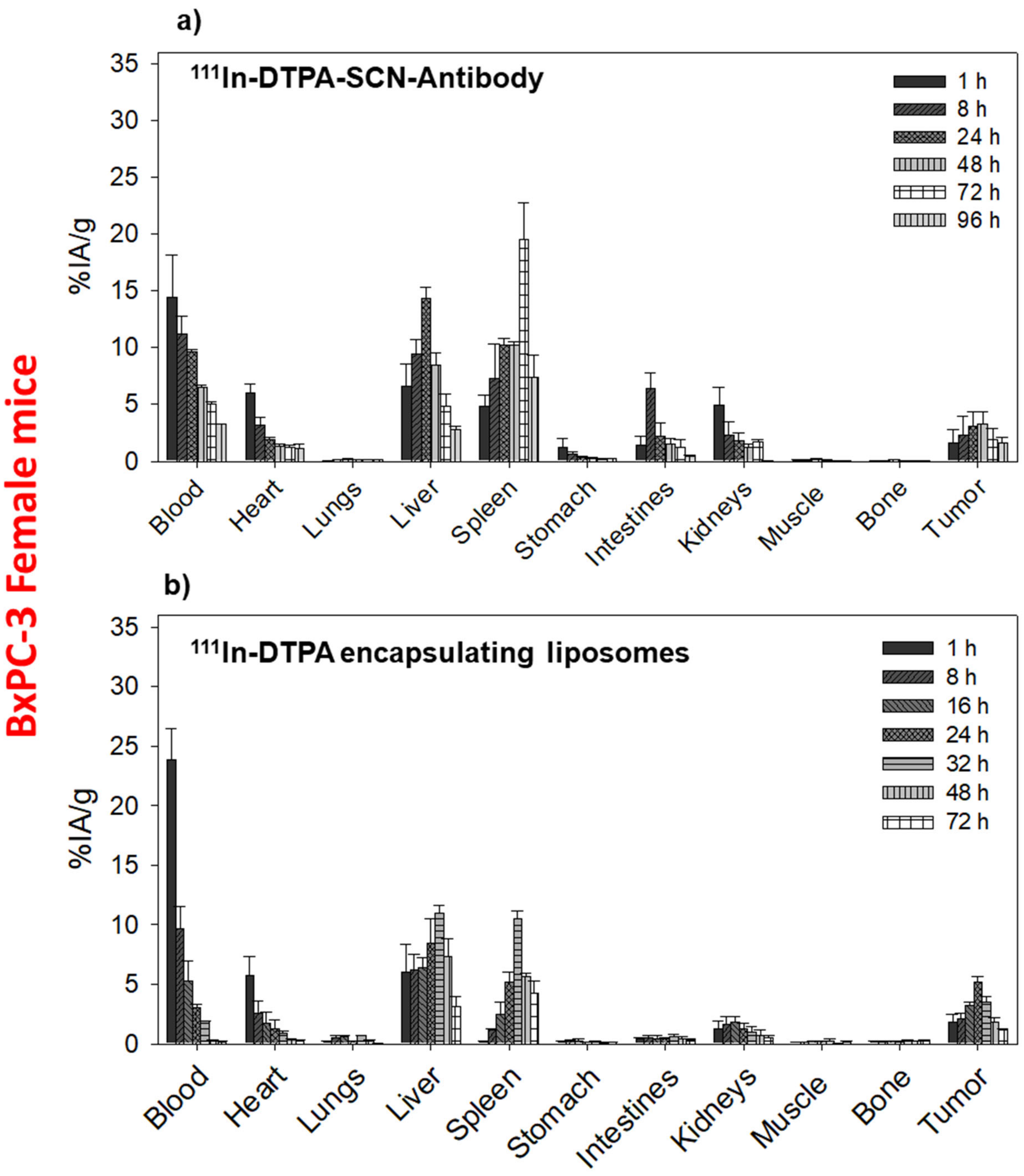

**Figure S12**. Biodistributions of (a) the HER1-targeting $^{111}$In-DTPA-SCN-antibody, Cetuximab, and the (b) tumor-responsive liposomes loaded with $^{111}$In-DTPA on **NSG male mice bearing BxPC3** tumors. $^{111}$In was used as surrogate of the parent $^{225}$Ac. Error bars correspond to standard deviations of n=3 mice per condition per time point.

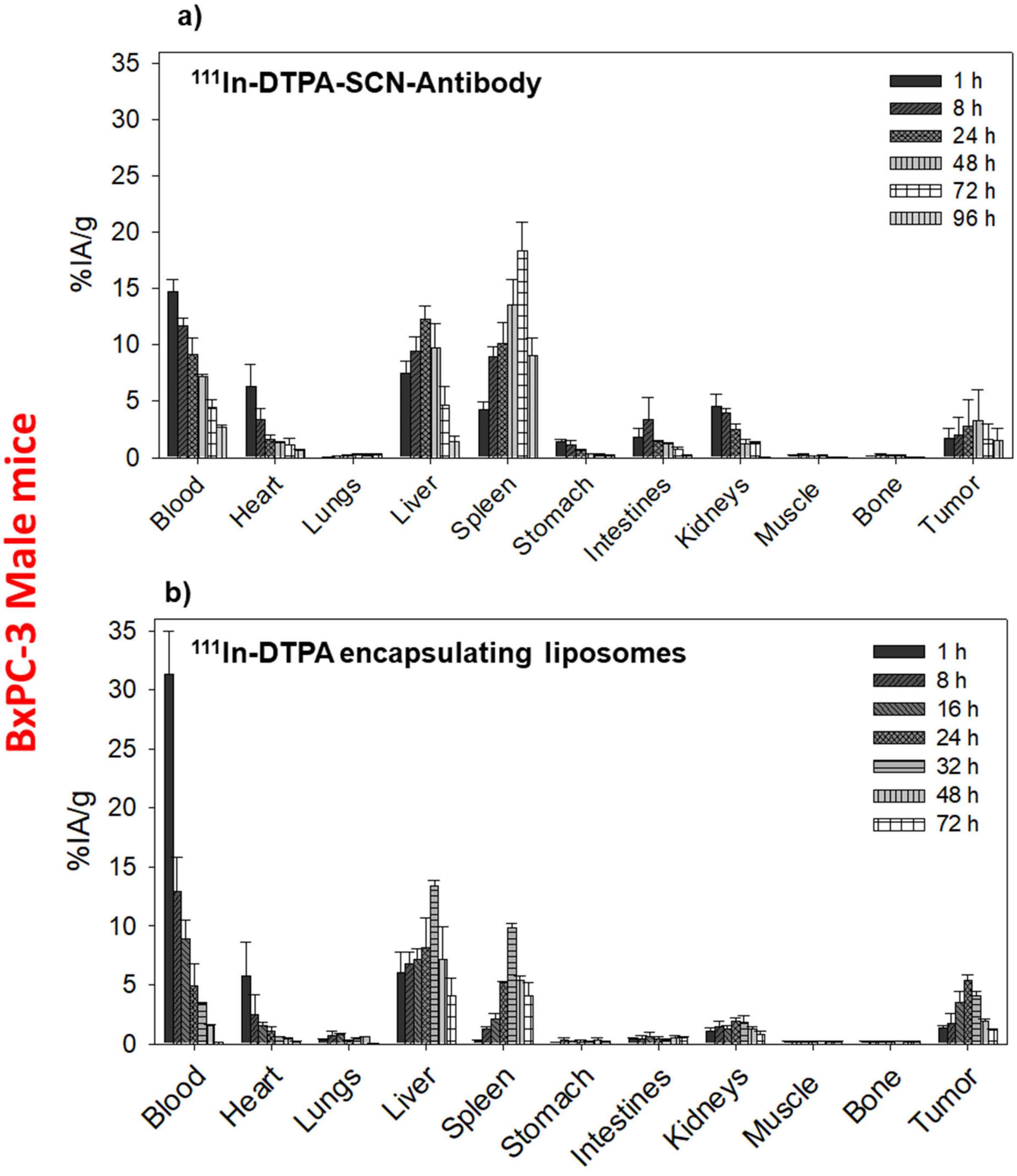

**Figure S13**. Tumor (top panel) and blood (lower panel) uptake- and clearance-kinetics of the HER1-targeting $^{111}$In-DTPA-SCN-Cetuximab (white symbols), and the tumor-responsive liposomes loaded with $^{111}$In-DTPA (black symbols) on NSG mice bearing tumors with HER1-expression levels varying as follows: CR-MDA-MB-436 > MDA-MB-231 > BxPC-3. $^{111}$In was used as surrogate of the parent $^{225}$Ac. The tumor delivered dose by the HER1-targeting antibody decreased with decreasing HER1-expression on tumor cells (left to right, top panel). The blood clearance kinetics for each carrier were comparable across all animal models (lower panel). Error bars correspond to standard deviations of n=3 mice per condition per time point.

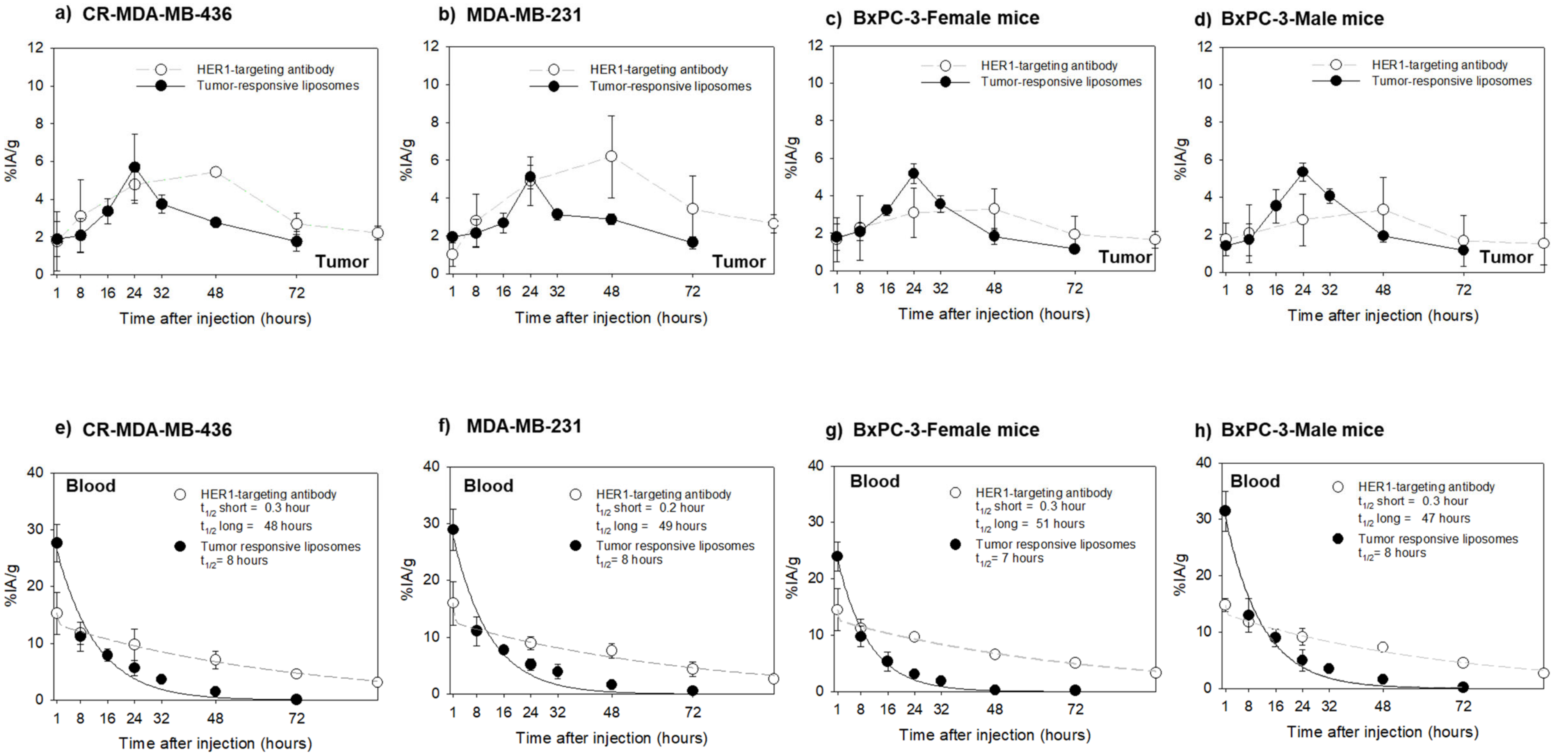

**Figure S14**. Volume of individual **CR-MDA-MB-436** primary, orthotopic, tumors over time following treatment with a single i.v. injection of 2.96 kBq $^{225}$Ac per 20 g mouse delivered by the HER1-targeting Cetuximab, the tumor-responsive liposomes, and/or both, separate, carriers at split ratios of radioactivity as indicated on plots.

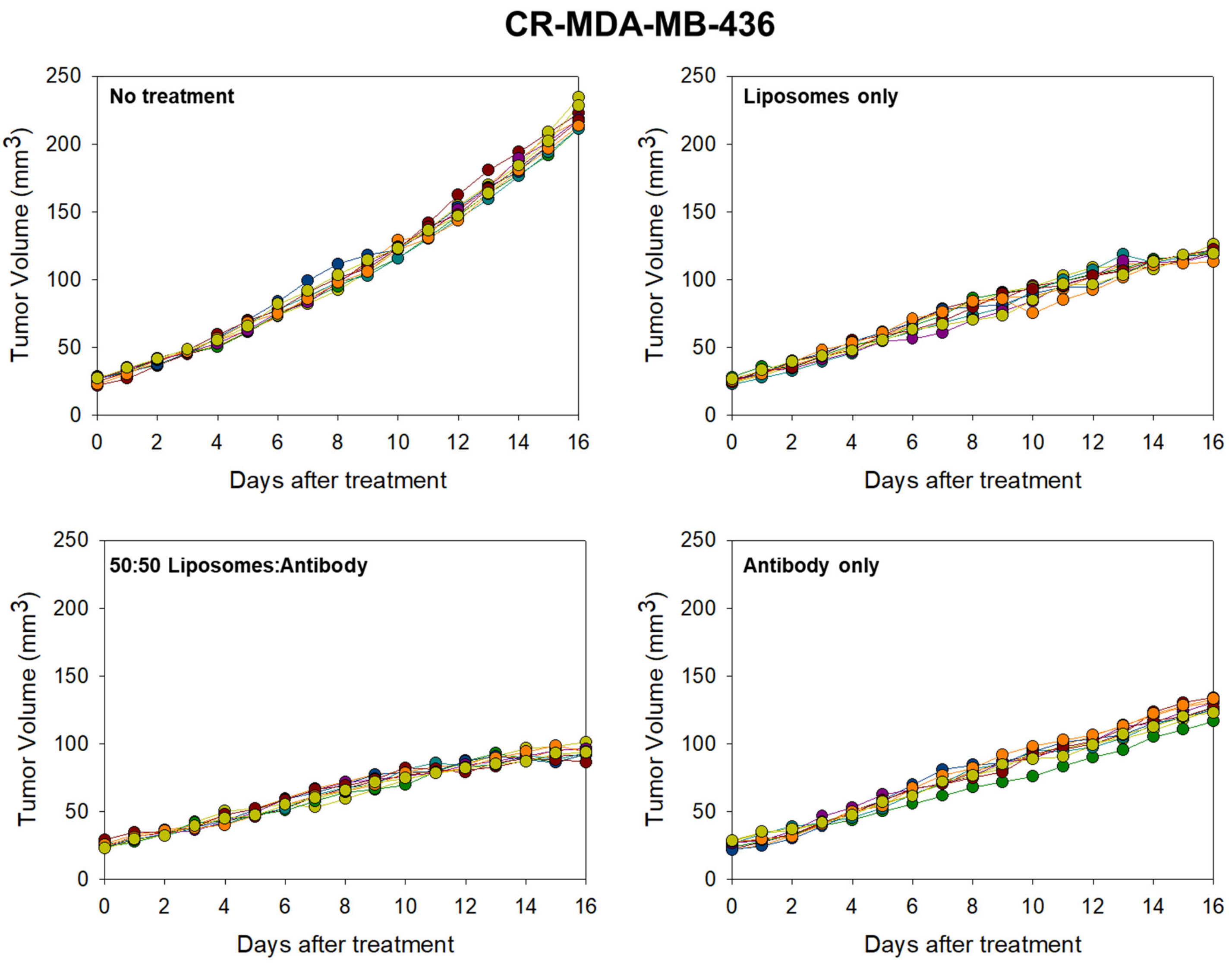

**Figure S15**. Volume of individual **MDA-MB-231** primary, orthotopic, tumors over time following treatment with a single i.v. injection of 2.96 kBq $^{225}$Ac per 20 g mouse delivered by the HER1-targeting Cetuximab, the tumor-responsive liposomes, and/or both, separate, carriers at split ratios of radioactivity as indicated on plots.

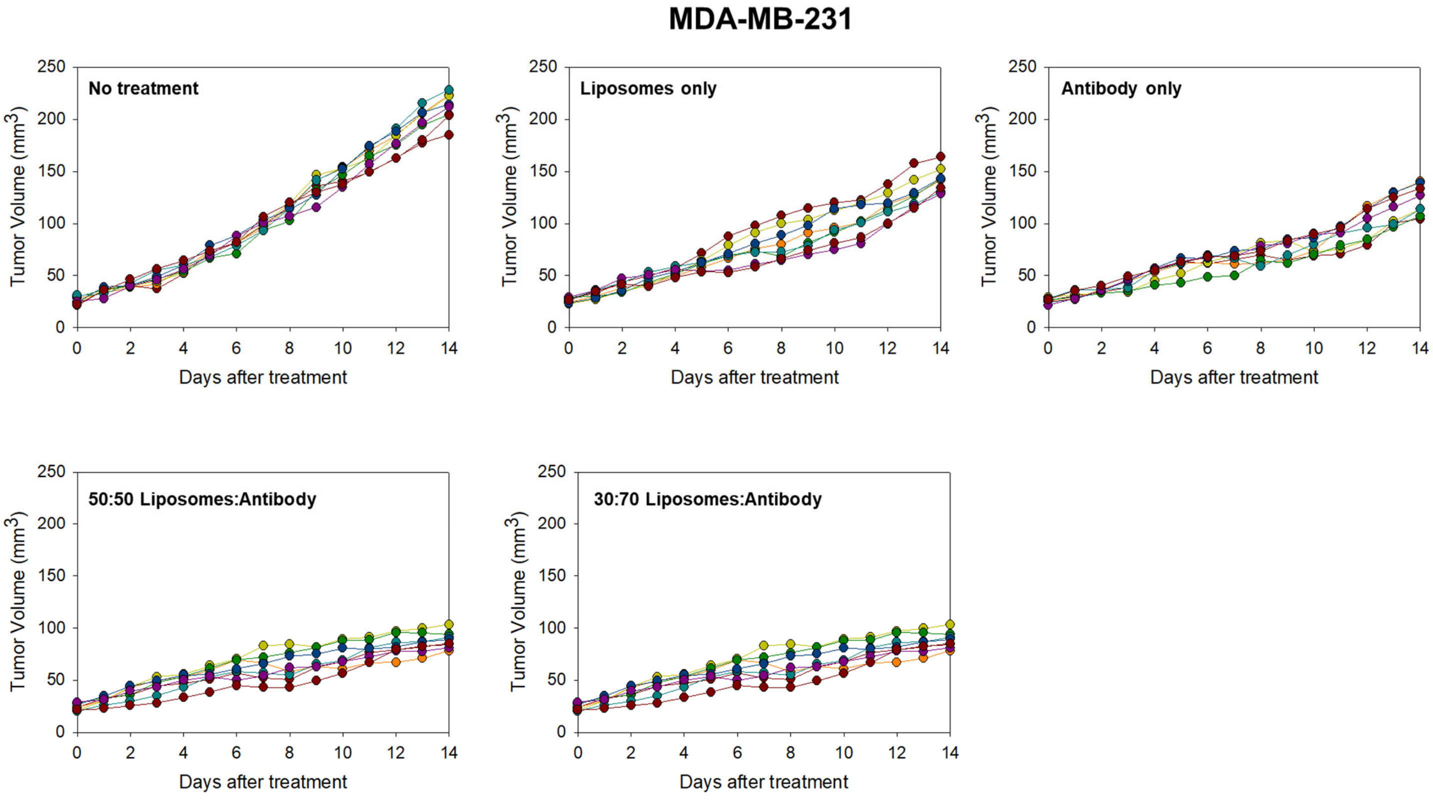

**Figure S16**. Volume of individual **BxP-3** subcutaneous tumors over time on NSG **female mice** following treatment with a single i.v. injection of 2.96 kBq $^{225}$Ac per 20 g mouse delivered by the HER1-targeting Cetuximab, the tumor-responsive liposomes, and/or both, separate, carriers at split ratios of radioactivity as indicated on plots.

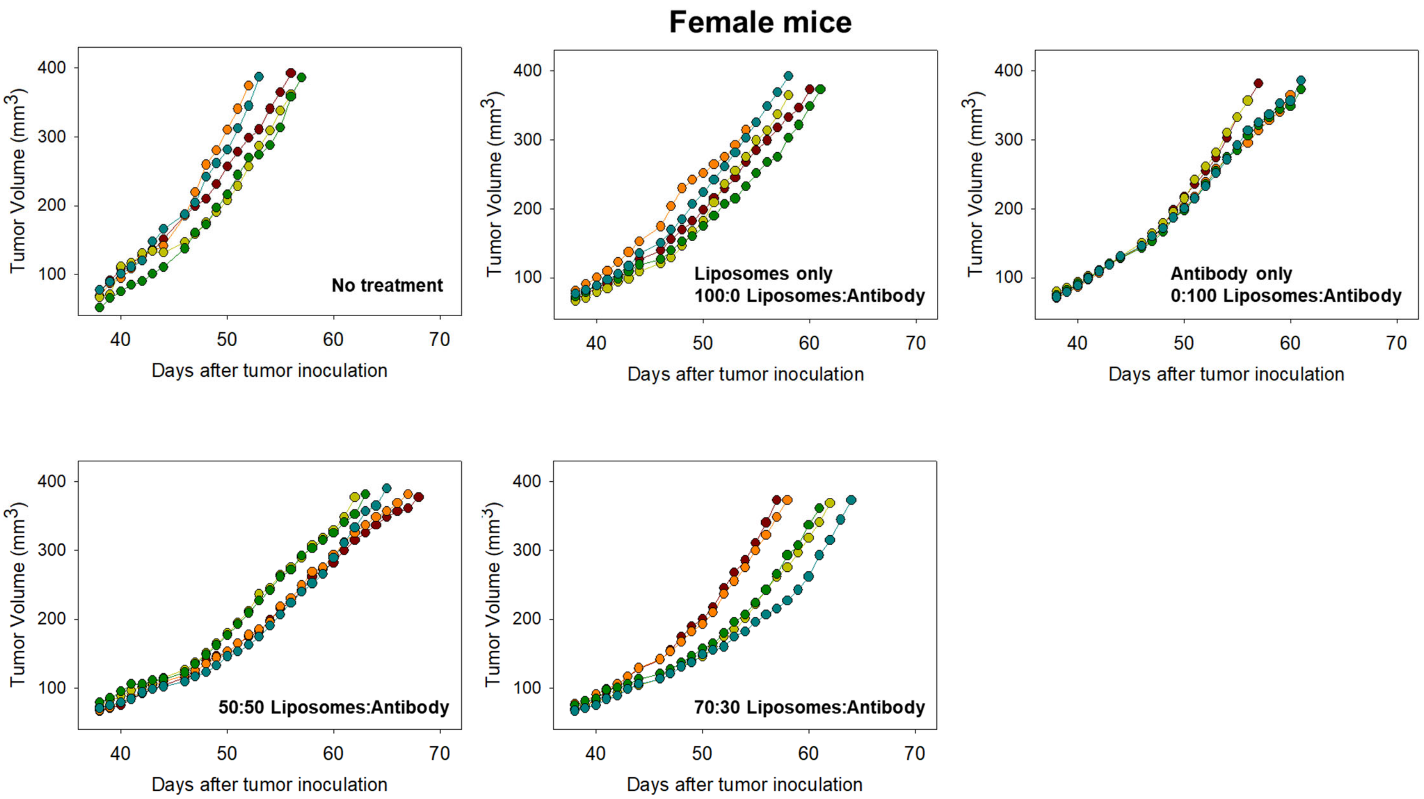

**Figure S17**. Volume of individual **BxP-3** subcutaneous tumors over time on NSG **male mice** following treatment with a single i.v. injection of 2.96 kBq $^{225}$Ac per 20 g mouse delivered by the HER1-targeting Cetuximab, the tumor-responsive liposomes, and/or both, separate, carriers at split ratios of radioactivity as indicated on plots.

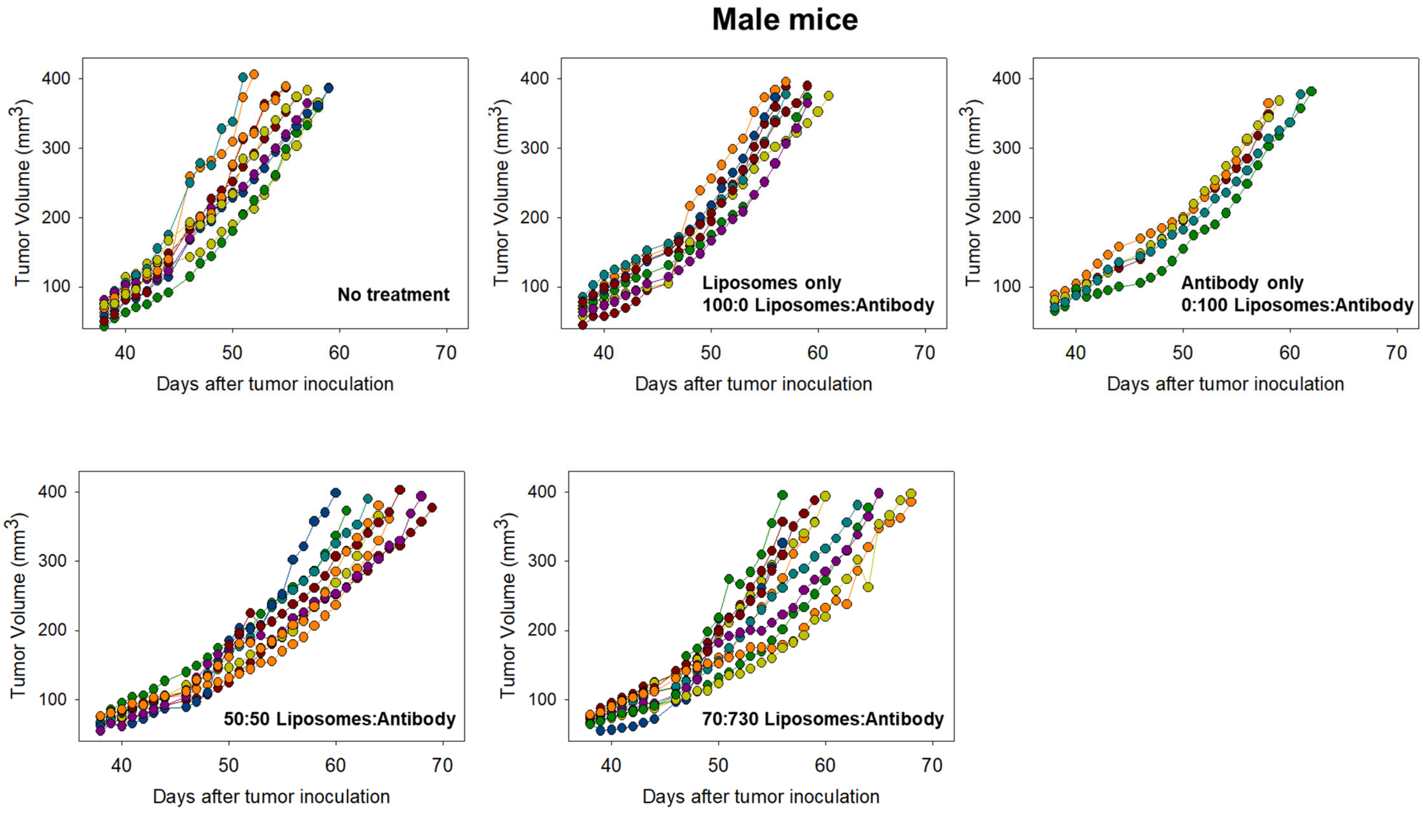

**Figure S18**. **Characteristic H&E stained sections of CR-MDA-MB-436 orthotopic tumors and of normal organs** from each treatment group, shown on Figure 3, acquired at the endpoint of the study. Scale bar=100μm.

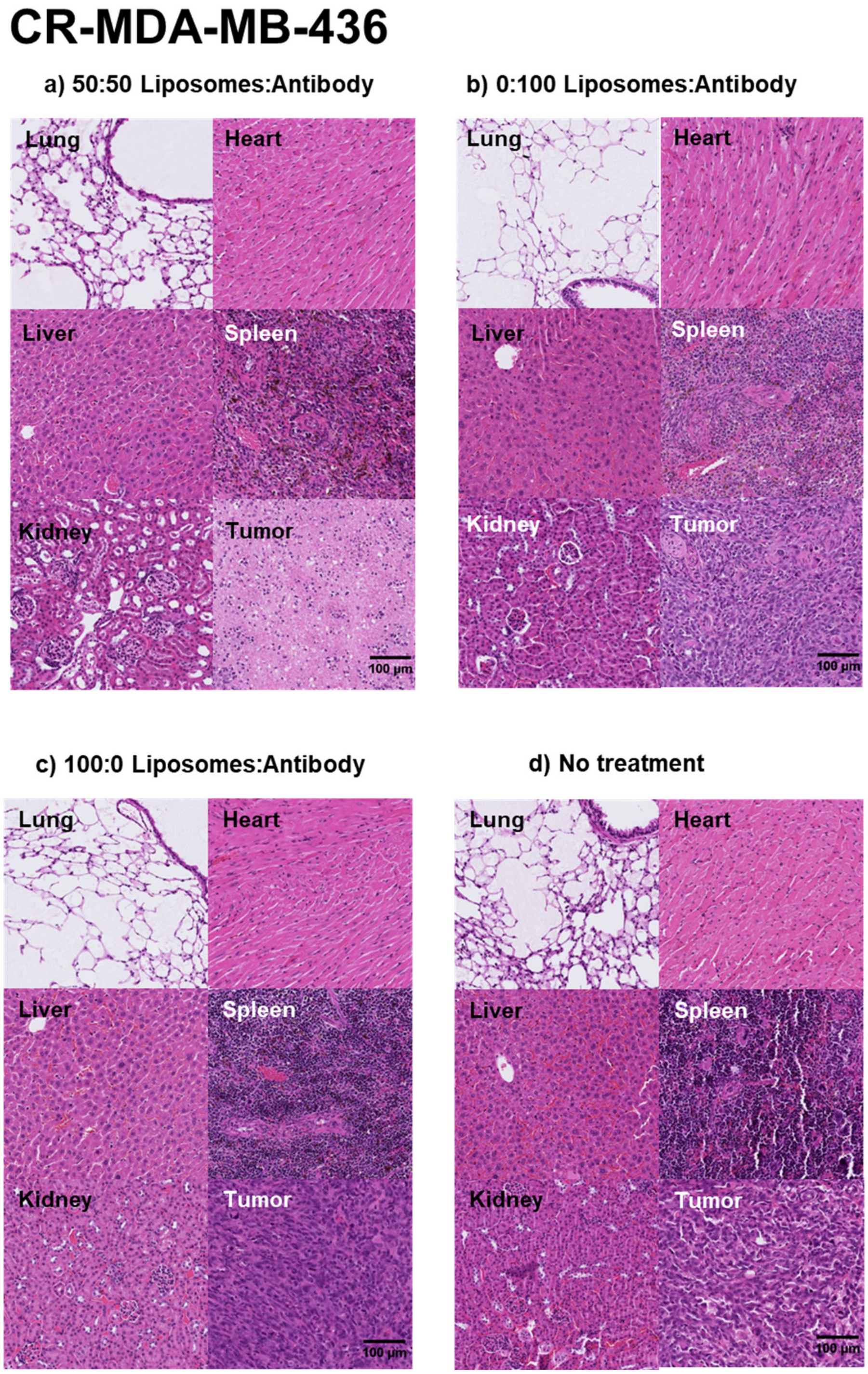

**Figure S19**. **Characteristic H&E stained sections of MDA-MB-231 orthotopic tumors and of normal organs** from each treatment group, shown on Figure 4, acquired at the endpoint of the study. Scale bar=100μm.

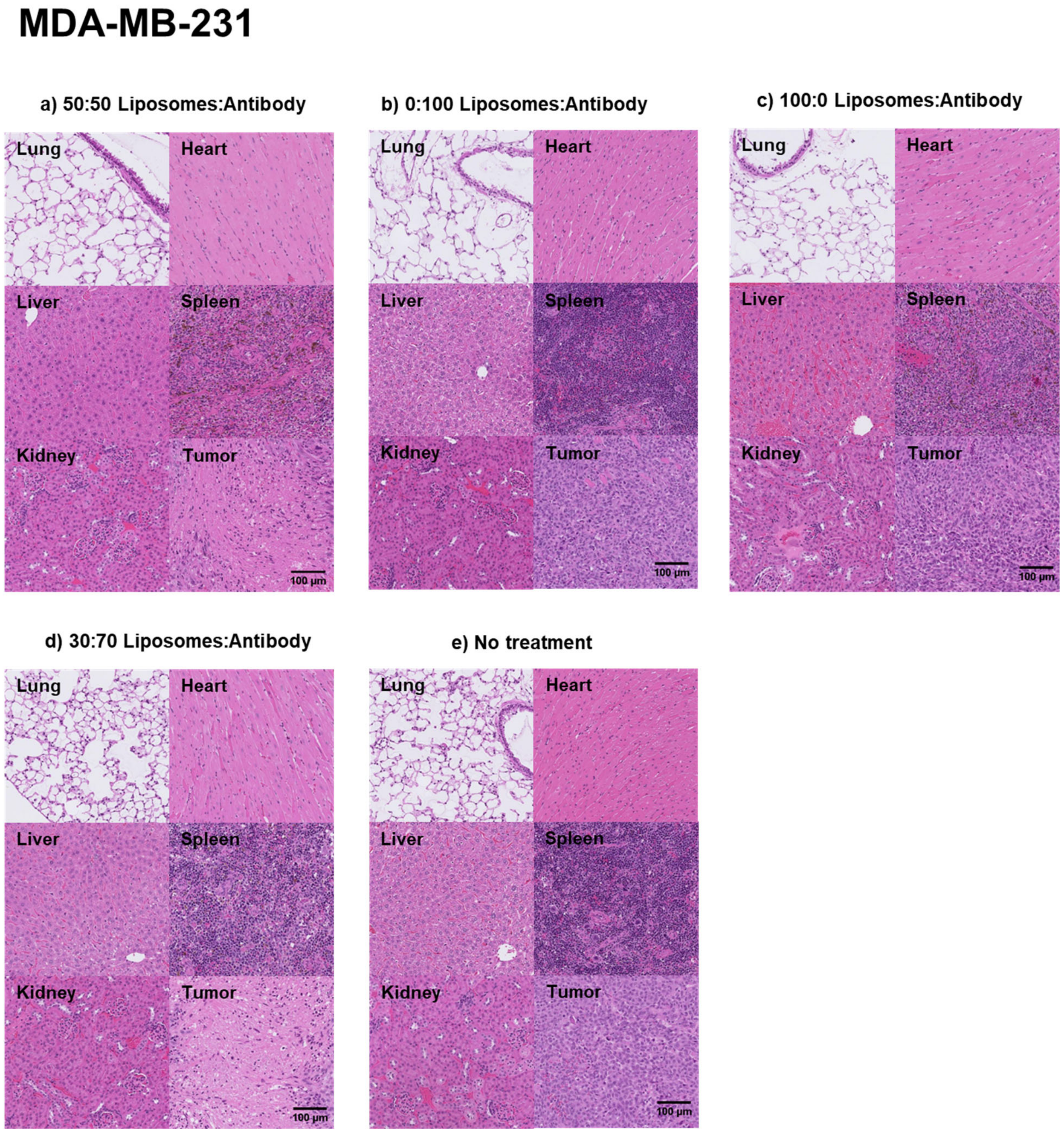

**Figure S20**. **Animal weights** over time during the treatment study on the **CR-MDA-MB-436** orthotopic mouse model, shown on Figure 3.

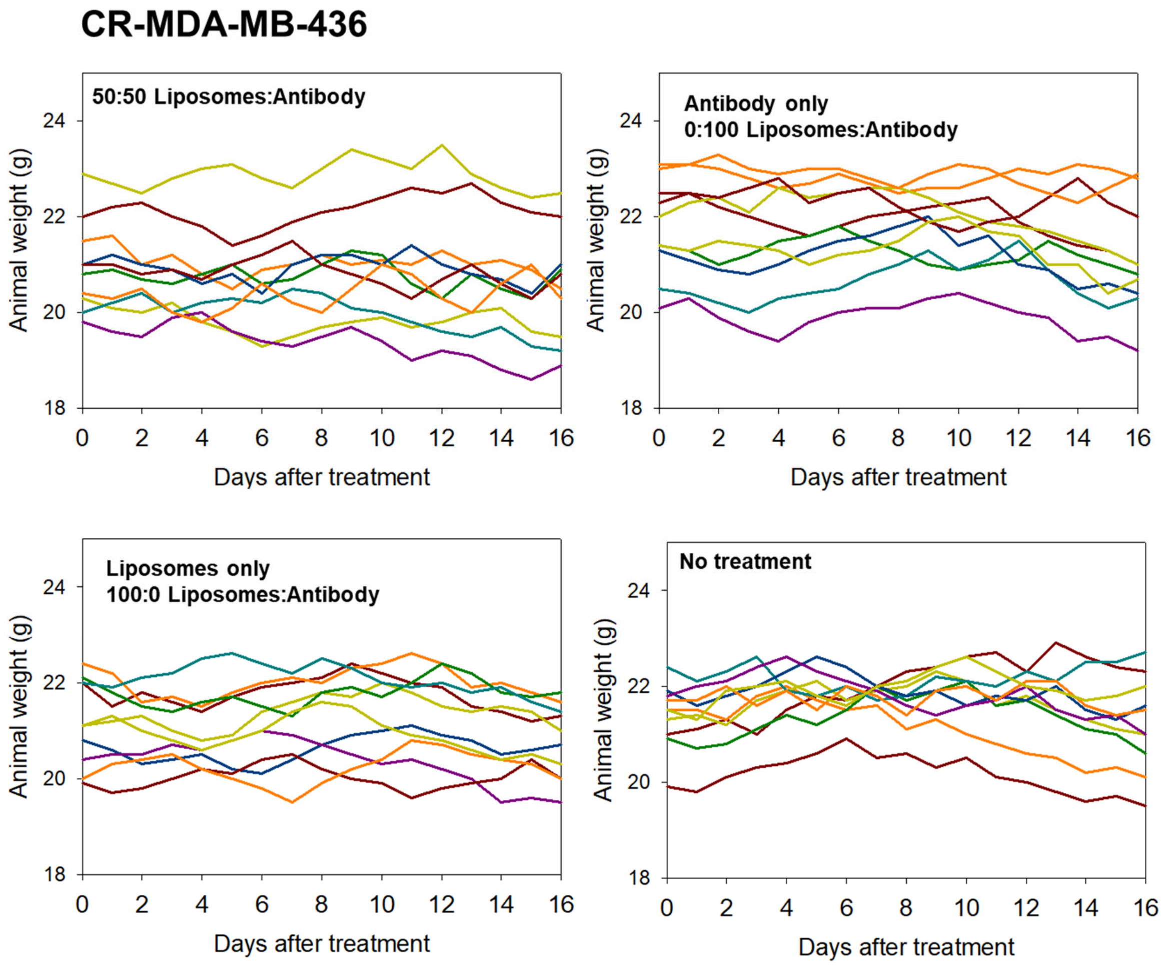

**Figure S21**. **Animal weights** over time during the treatment study on the **MDA-MB-231** orthotopic mouse model, shown on Figure 4.

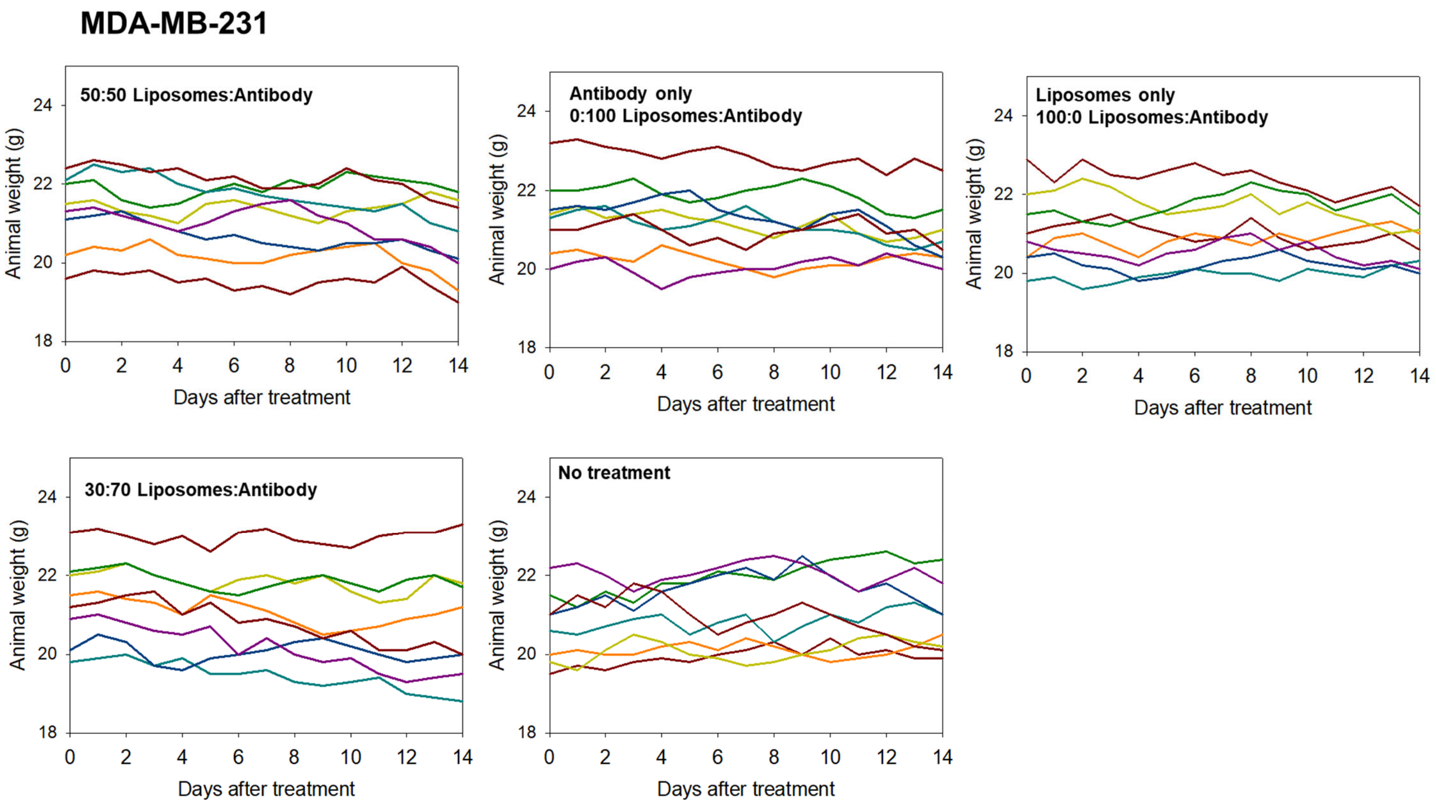

**Figure S22**. **Characteristic H&E stained sections of BxPC3 subcutaneous tumors and of normal organs** from each treatment group on female and male mice, shown on Figure 5, acquired at the endpoint of the study. Scale bar=100μm.

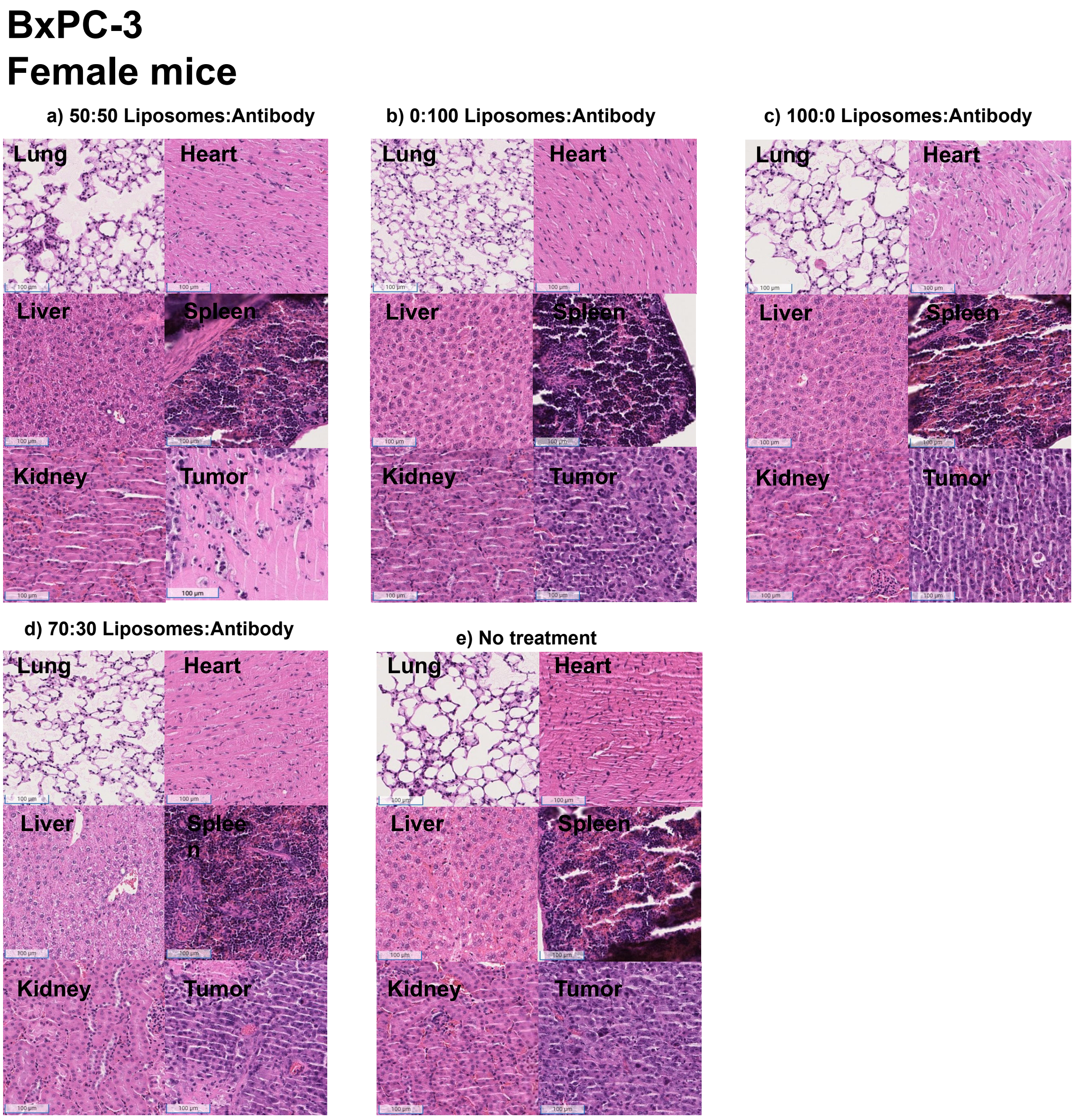

### BxPC-3
### Male mice

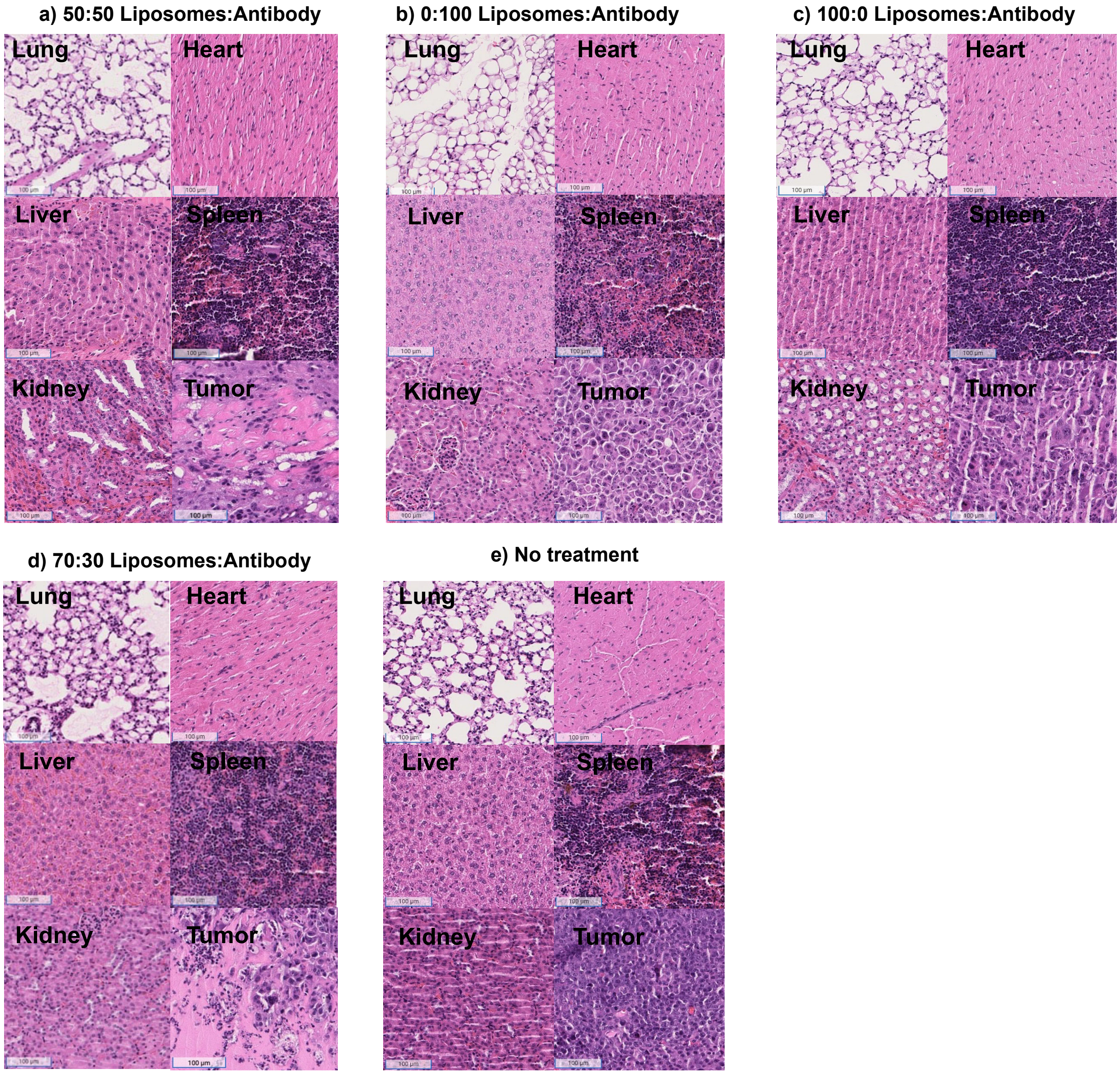

**Figure S23**. **Characteristic H&E stained sections of BxPC3 subcutaneous tumors on male mice treated with two injections of therapy,** shown on Figure 6, acquired at the endpoint of the study. Scale bar=100μm

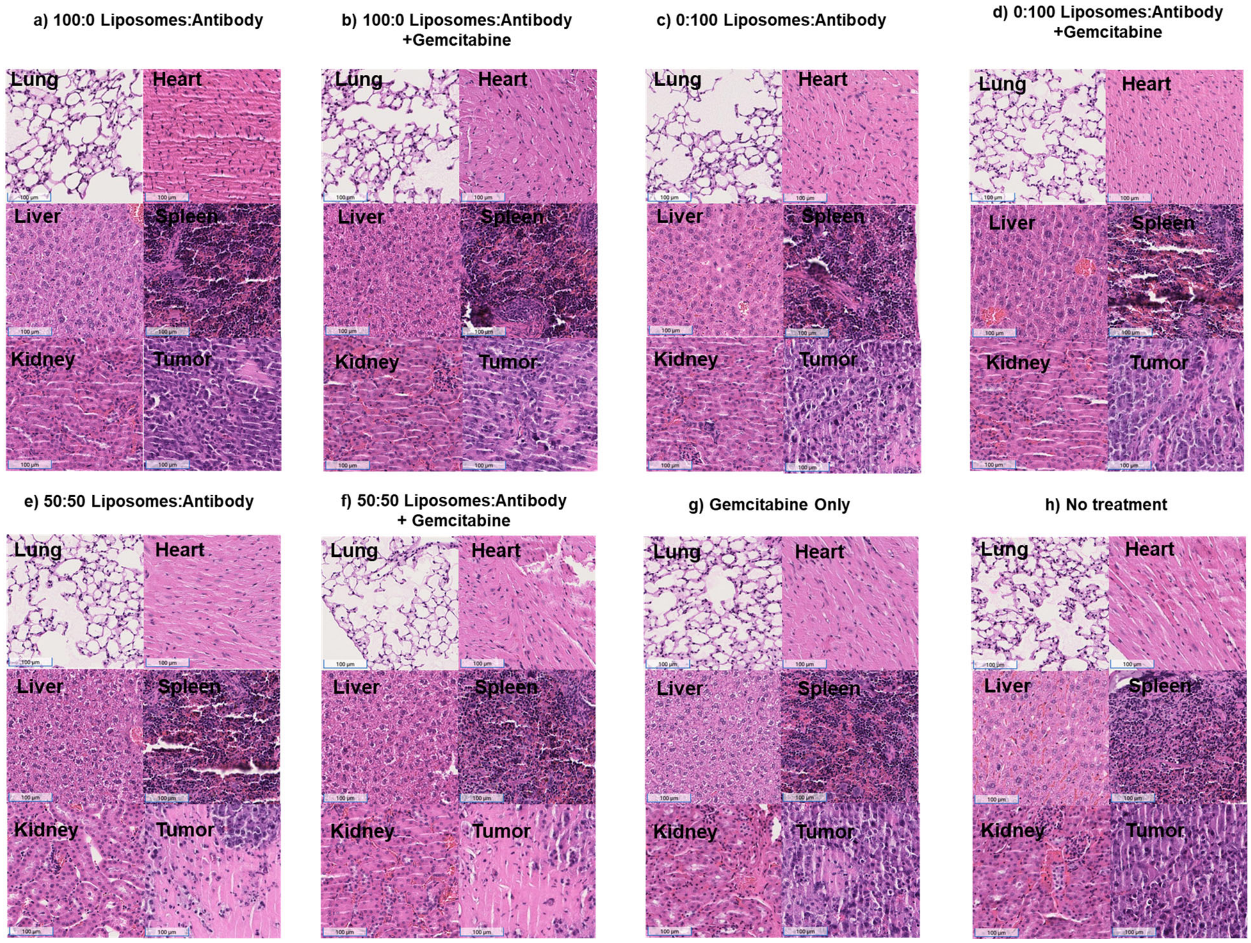

**Figure S24**. **Animal weights** over time during the treatment study on the **BxPC3** subcutaneous female and male mouse models, shown on Figure 5.

**BxPC-3**
**Female mice**

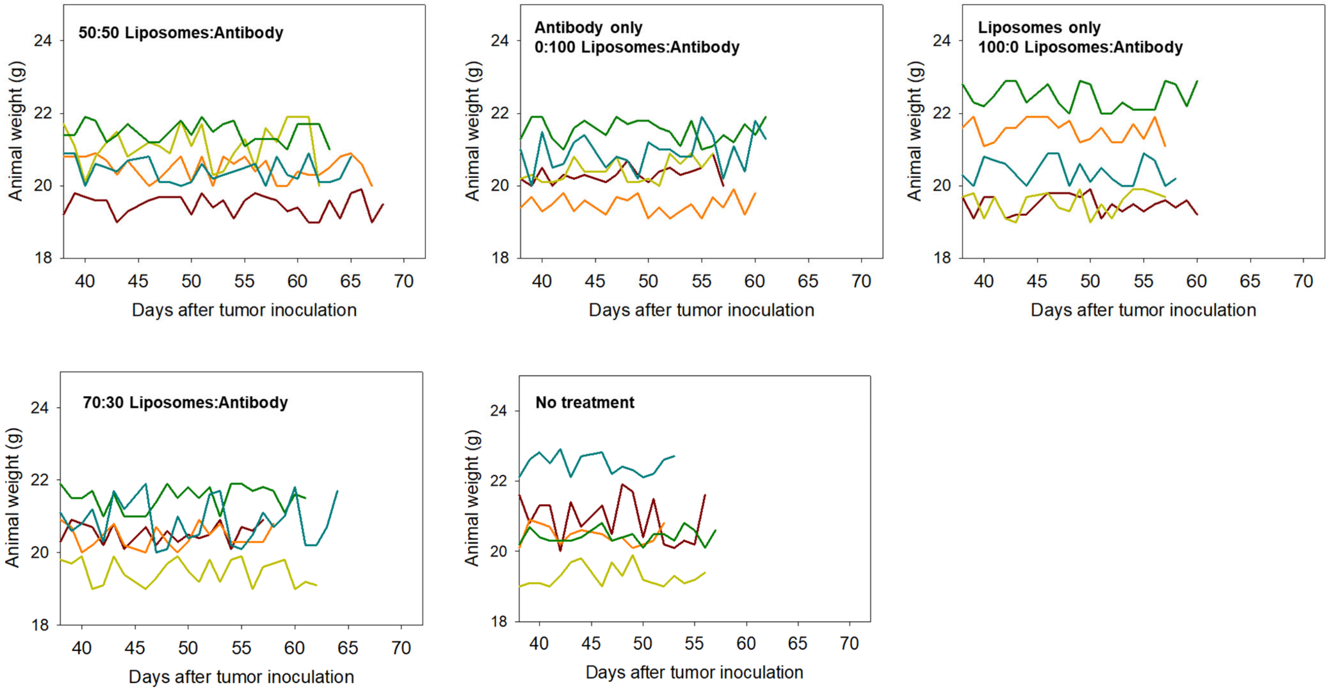


**BxPC-3**
**Male mice**

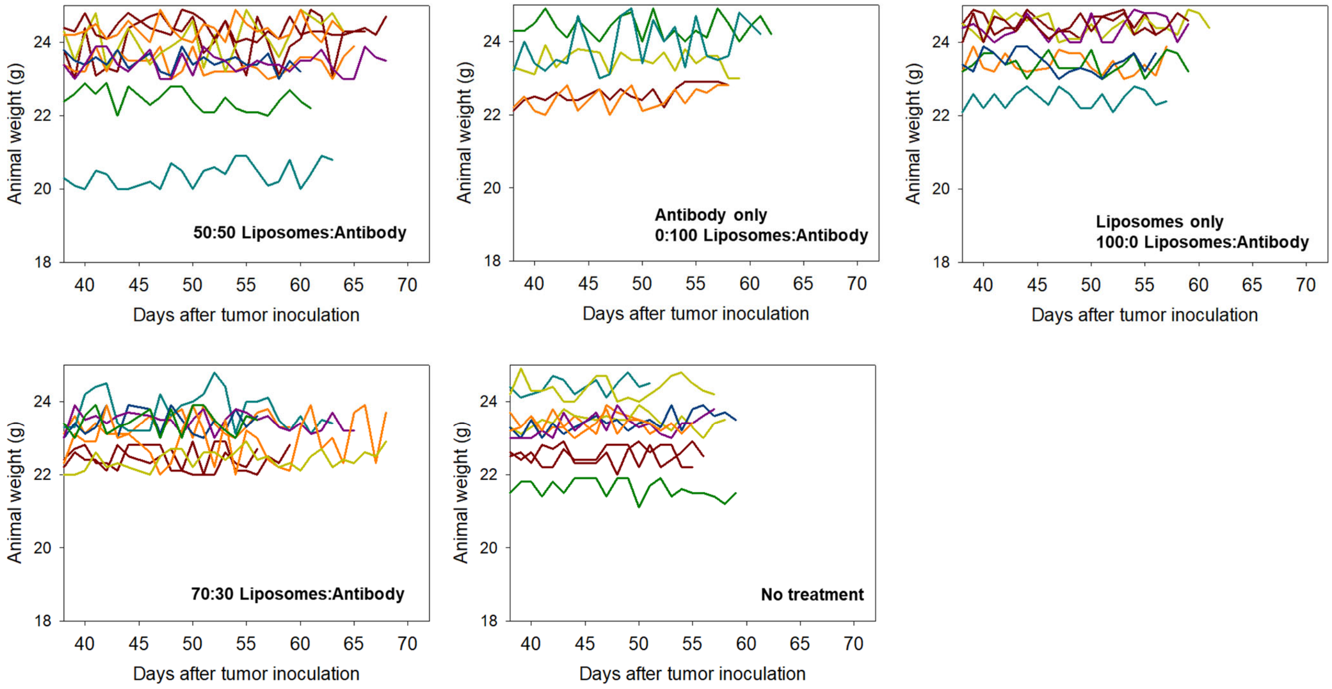

**Figure S25**. **Animal weights** over time during the treatment study on the **BxPC3** subcutaneous male mouse model, shown on Figure 6.

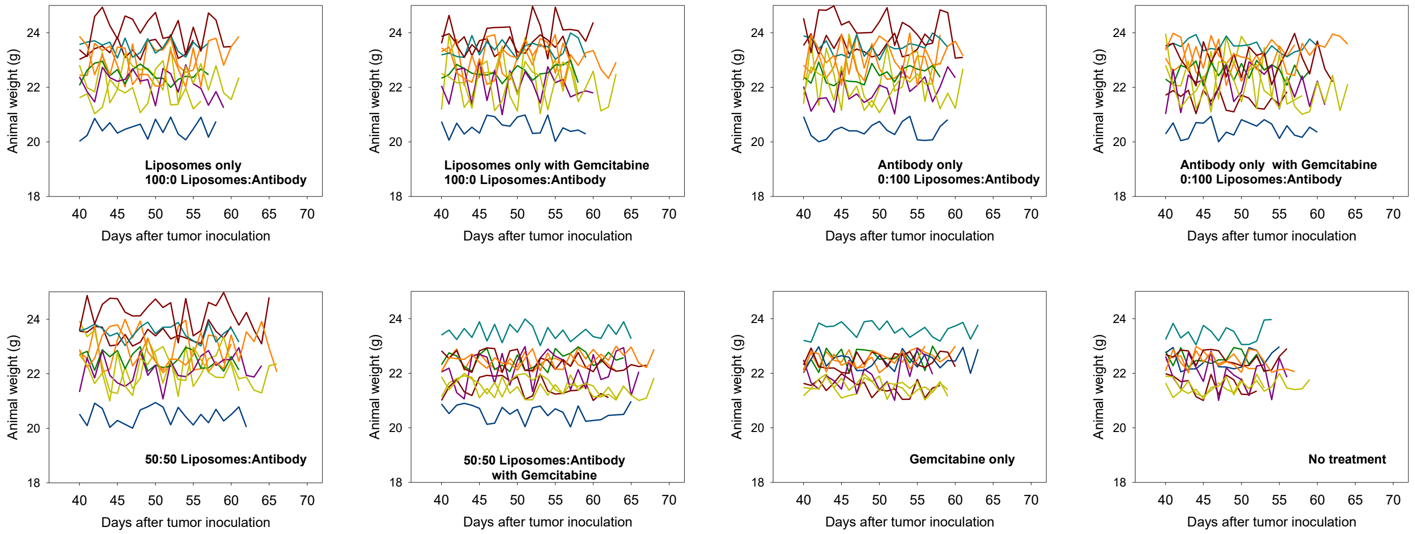

**Figure S26**. On liposomes, the releasing property (R) is key in enabling the highly diffusing $^{225}$Ac-DOTA to irradiate the areas of spheroids that are far from the edge (3). **Outgrowth inhibition of 400 μm in diameter MDA-MB-231 spheroids upon treatment with combinations of non-releasing (R-, left) or releasing (R+, right) liposomes loaded with $^{225}$Ac-DOTA and the HER1-targeting $^{225}$Ac-DOTA-SCN-Cetuximab.** The total radioactivity concentration was kept constant at 3.7 kBq/mL. Error bars correspond to standard deviations of repeated measurements (n=12 spheroids per condition, n=2 independent liposome and antibody preparations). * indicates *p*-values < 0.05.

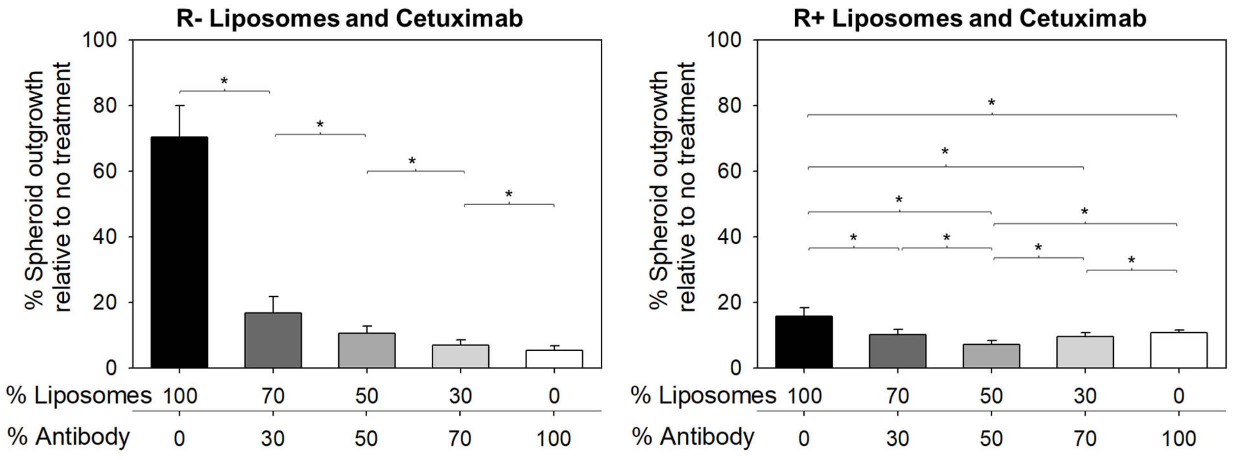
